# Assessing the Extent of Structural and Dynamic Modulation of Membrane Lipids due to Pore Forming Toxins: Insights from Molecular Dynamics Simulations

**DOI:** 10.1101/2020.01.13.905463

**Authors:** Vadhana Varadarajan, Rajat Desikan, K. G. Ayappa

## Abstract

Infections in many virulent bacterial strains are triggered by the release of pore forming toxins (PFTs), which form oligomeric transmembrane pore complexes on the target plasma membrane. The spatial extent of the perturbation to the surrounding lipids during pore formation is relatively unexplored. Using all-atom molecular dynamics simulations, we investigate the changes in the structure and dynamics of lipids in a 1,2-*dimyristoyl-sn-glycero-3-phosphocholine* (DMPC) lipid bilayer in the presence of contrasting PFTs. Cytolysin A (ClyA) an *α* toxin with its inserted wedge shaped bundle of inserted *α* helices induces significant asymmetry across the membrane leaflets in comparison with *α* hemolysin (AHL) a *β* toxin. Despite the differences in hydrophobic mismatch and uniquely different topologies of the two oligomers, perturbation to lipid order as reflected in the tilt angle and order parameters, and membrane thinning is short ranged, lying within ∼ 2.5 nm from the periphery of the either pore complex, commensurate with distances typically associated with van der Waals forces. In contrast, the spatial extent of perturbations to the lipid dynamics extend outward to at least 4 nm for both proteins, and the continuous survival probabilities reveal the presence of a tightly bound shell of lipids in this region. Displacement probability distributions show long tails and the distinctly non-Gaussian features reflect the induced dynamic heterogeneity. A detailed profiling of the protein-lipid contacts with residues tyrosine, tryptophan, lysine and arginine show increased non-polar contacts in the cytoplasmic leaflet for both PFTs, with a higher number of atomic contacts in the case of AHL in the extracellular leaflet due to the mushroom-like topology of the pore complex. The short ranged nature of the perturbations observed in this simple one component membrane suggests an inherent plasticity of membrane lipids enabling recovery of structure and membrane fluidity even in the presence of these large oligomeric trans-membrane protein assemblies. This observation has implications in membrane repair processes such as budding or vesicle fusion events used to mitigate PFT virulence, where the underlying lipid dynamics and fluidity in the vicinity of the pore complex are expected to play an important role.

## 1 Introduction

The cell membrane is a highly heterogeneous and crowded semi-permeable barrier, within which, membrane-inserted proteins can exist in a variety of complex dynamic oligomeric architectures implicated in cell signalling and transport. ^1^ Unravelling the mechanisms and energetics of the insertion of transmembrane proteins, formation of intricate membrane-inserted architectures, their function, and their influence on the surrounding lipid milieu is a challenging problem in membrane biophysics. Large oligomeric transmembrane architectures not native to the plasma membrane are formed by several toxic proteins whose primary aim is to disable the functioning of the cell. ^2^ In this regard, bacterial pore-forming toxins (PFTs) represent a large class of proteins which play an important role in bacterial pathogenesis implicated in many virulent diseases such as cholera, anthrax, pneumonia and listeriosis. PFTs are initially expressed as water soluble monomers, which upon binding to the cell membrane spontaneously self-assemble to form transmembrane pore complexes with large extracellular solvent-exposed domains. The PFTs are broadly classified as *α* and *β* toxins depending on the secondary structure of the transmembrane domains, which are the either *α*-helices for the former or *β*-barrels for the latter. Unregulated pore formation leads to osmotic imbalance and eventually cell lysis. Unlike certain families of PFTs which require specific receptors or the presence of cholesterol in the membrane, several PFTs such as Cytolysin A (ClyA) an *α*-PFT and *α*-hemolysin (AHL) a *β*-PFT have evolved to form pores in a wide range of membrane environments. This unique oligomerization ability is unlike transmembrane proteins such as serotonin receptors for example, whose functioning is intrinsically connected to the underlying membrane composition ^3,4^.

In order to defend the cell against attack by PFTs, several repair pathways which involve removal of the membrane regions where PFT oligomers are present have evolved. ^5^ Among these pathways exocytosis, ectosytosis and endocytosis ^6,7^ involve large scale membrane re-organization such as vesicle fusion or budding events where the underlying lipid dynamics and fluidity play an important role. For example during ectocytosis, and vesicle shedding, budding occurs around the region containing the pore forming oligomers and it is essential for the underlying membrane to have sufficient fluidity to export lipids during bud formation leading to eventual pinch-off to clear the protein bound sites on the membrane. While investigating PFT action on model biomembrane systems, the focus is largely on the influence of lipid composition on pore formation pathways and kinetics, ^8,9^ the ensuing lipid rearrangement and dynamics, challenging to capture experimentally, has received less attention. Hence the the spatial extent to which a fully formed PFT oligomer perturbs the surrounding lipids both structurally and dynamically is not completely understood. Models have invoked the notion of an effective hydrodynamic radius for proteins which includes the protein along with a strongly bound shell of lipids to reconcile data from photobleaching experiments. ^10,11^ Whether a similar situation exists with the more complex oligomeric PFTs is not known.

Understanding the role and rearrangement of lipids in proteolipidic complexes is only recently getting attention due to advances in nuclear magnetic resonance (NMR) techniques where the active role of lipids as regulators to protein function and membrane re-organization is being unravelled. ^12^ The role of lipids in stabilizing transmembrane oligomers as a function of oligomerization has been recently illustrated using mass spectrometry. ^13^ Proteins can adopt different orientations to minimize hydrophobic mismatch ^14^ as well as induce changes in the underlying phases of the phospholipid membranes. ^15^

Multimeric pore assemblies formed from PFTs range from 7 − 12 monomeric units for the smaller pore complexes formed by cytolysin A and *α*-hemolysin, to 30 − 40 units in the case of cholesterol dependent cytolysins such as listeriolysin O (LLO) and pneumolysin. ^2^ In contrast to native transmembrane proteins, PFT architectures predominantly consist of smaller transmembrane segments and larger extracellular domains which can peripherally interact with the extracellular membrane leaflet. ^2^ This unique architecture results in asymmetric interactions with the lipids, leading to curvature induced membrane stresses. With the advent of super-resolution techniques such as stimulated emission depletion coupled with fluorescence correlation spectroscopy (STED-FCS) lipid dynamics in cells and supported membrane platforms can be resolved at length scales below 100 nm. ^16–20^ STED-FCS experiments capture a cross-over at smaller length scales (< 200 nm) from Brownian to non-Brownian dynamics with increasing cholesterol content in supported lipid bilayers exposed to LLO a cholesterol dependent cytolysin ^18^ as well as increased fluidization in three component raft forming membranes upon ClyA binding. ^21^ In a recent study with listeriolysin O, the dynamics of lipids on giant unilamellar vesicles were found to be intrinsically coupled to the state of the membrane bound protein. ^22^

The role of lipids and their interactions with transmembrane proteins have also been investigated using MD simulations. For example, Woolf and Roux ^23^ in a pioneering molecular dynamics study of the gramicidin A channel inserted into a *1-2, dimyristoylsn-glycero-3-phosphocholine* (DMPC) bilayer (< 1 ns) illustrate the ordering and interaction energetics of lipids with specific protein residues such as tryptophan and leucine. In a comparative study, Sansom ^24^ and co-workers contrasted protein-lipid interactions of the pottasium channel KcsA with it’s membrane inserted *α* helices with OmpA a *β*-barrel protein using 20 ns long MD simulations. Mobility differences between bound and free lipids were discerned from these simulations and a lowered mobility was observed for lipids in the vicinity of the KcsA protein. More recently 30*µ*s coarse grained simulations with over 60 different lipid types have been used to illustrate the preferential segregation of lipids around different classes of integral membrane proteins. ^25^

In this manuscript, we investigate the perturbation and heterogeneity induced in 1,2-*dimyristoyl-sn-glycero-3-phosphocholine* (DMPC) lipids in the presence of transmembrane pores of ClyA and AHL. The transmembrane pores formed by each of these PFT’s are illustrated in Fig. 1, and some distinct differences emerge. The ClyA pore consists of an *α*-barrel extracellular domain with outer and inner diameters of 10.5 and 7 nm respectively, and membrane inserted amphiphatic *α*-helices, which taper into a wedge-shaped domain in the cytoplasmic leaflet (CL) with reduced outer (9 nm) and inner (4 nm) diameters. The membrane inserted structure for AHL is distinctly different with a transmembrane region comprises of a 2.8 nm wide *β*-barrel and a cap-like region which engages in extended lateral interactions with the extra-cellular leaflet (EL). One of the main goals of this work is to study the spatial extent of structural and dynamic heterogeneities induced to the surrounding lipids due to the presence of these topologically and distinct classes of pore forming toxins. All atom molecular dynamics simulations (0.5*µ*s) of both the pore complexes in a DMPC bilayer are carried out to compare and contrast the lipid order parameters, bilayer thickness, leaflet specific lipid dynamics, lifetime analysis, hydrogen bonding patterns and survival probabilities. In addition to being representative of a phosphatidylcholine (PC) lipid, DMPC results in a hydrophobic mismatch only in the case of ClyA, allowing us to contrast the extent to which this influences the spatial modulation in the surrounding lipids. We analyze and partition interactions of the lipids with protein residues such as arginine, lysine, tryptophan and tyrosine that have been implicated in electrostatic and van der Waals protein-lipid interactions. Despite specific differences in the lipid structure and dynamics, the spatial extent of structural modulation in the lipids is short ranged when compared with the longer ranged dynamical modulation. Interestingly the range of such spatial perturbations is similar for both classes of PFTs.

**Fig. 1.**
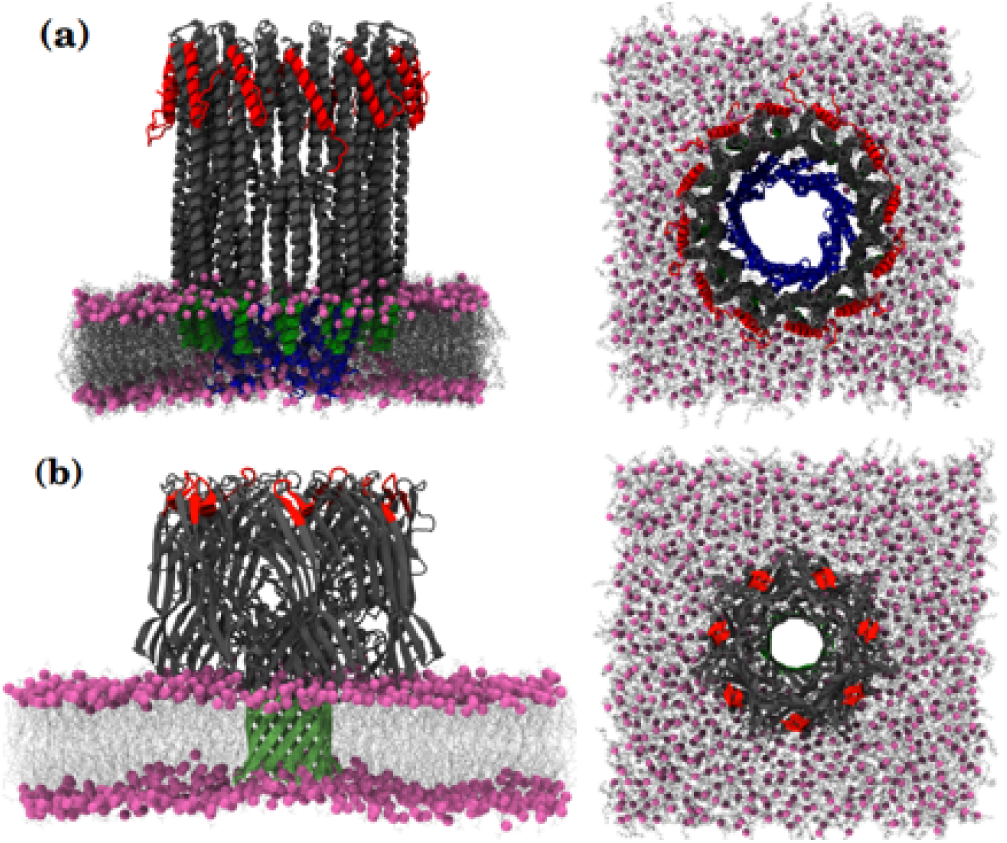
Sideviews of a lipid bilayer embedded with pore forming toxins are illustrated on the left with corresponding topviews on the right. The phosphorous head groups in the bilayer are represented in mauve. (a) ClyA, an *α*-PFT forms a dodecameric pore with *α* helices in the transmembrane region of the bilayer. The pore diameter is larger in the extracellular (EL) leaflet and tapers into a wedge shape at the cytoplasmic leaflet (CL). (b) AHL, a *β*-PFT forms a heptameric complex with a membrane inserted *β* barrel.

## 2 Computational Methods

### 2.1 Molecular modelling of the ClyA and AHL pores

The dodecameric crystal structure of the Cytolysin A (ClyA) pore without the ethyl mercury ion ligands was used as the starting structure (PDB ID 2WCD). ^26^ Disordered N-terminal residues 1-7 and the C-terminal residues 293-303 unresolved in the crystal structure are important for lytic activity ^27–29^ and hence were modeled using the I-TASSER web server ^30,31^ as described previously. ^29,32^ The heptameric crystal structure of the *α*-hemolysin (AHL) pore (PDB ID 7AHL) ^33^ has multiple residues with missing coordinates (arginine 66 and lysine 70 in chain A, lysine 30 and lysine 240 in chain D, lysine 283 in chain F and lysine 30 in chain G). The missing atoms in these residue side chains were reconstructed using VMD *1*.*9*.*1* ^34^ similar to a previous study. ^35^ MD simulations were performed using GROMACS version *4*.*6*.*4* (www.gromacs.org). ^36^ All the residues including histidine in both pores were set to default protonation states corresponding to a neutral pH with the pdb2gmx tool in GROMACS. Cartoon representations of both the ClyA and AHL pores is shown in Fig. 1, and all graphical rendering of simulation snapshots were performed with VMD.

### 2.2 Fully atomistic equilibrium MD simulations of the ClyA and AHL pores in DMPC membranes

The AMBER ff99SB-ILDN force-field with *φ* corrections ^37,38^ was used to describe the interactions of the protein atoms and the ions, along with the TIP3P water model ^39^ and the Amber force-field compatible ‘Slipid’ parameters for the saturated DMPC (1,2-dimyristoyl-sn-glycero-3-phosphocholine) lipids. ^40^ The modeled pore structures were inserted into pre-equilibrated DMPC membrane patches of size 18 nm*×*18 nm. All simulations were carried out at salt concentrations corresponding to a physiologically relevant 0.15 M NaCl. The number of lipids, water and salt molecules for the different simulations are given in Table S1 (ESI†) Electrostatic interactions were computed using the smooth Particle Mesh Ewald (PME) method ^41^ with a real space cut-off of 1.0 nm and a fourth order (cubic) interpolation. A 1.0 nm cut-off was used for computing the van der Waals. interactions, and these were shifted to a zero potential at the cut-off. All bonds were constrained using the LINCS algorithm. ^42^ Simulations were run using a leap-frog integrator with a 2 fs integration time step and with Verlet buffered lists (target energy drift of 0.005 kJ mol^-1^ ns^-1^ per atom). The neighbour list update frequency was once every 10 steps. Energy minimization after addition of solvent and ions with a displacement of 0.01 nm/step was carried out using the steepest descent method for 20000 steps. Short equilibration simulations for 50 ps in NVT and 500 ps in the NPT ensembles were performed with and without harmonic restraints on the protein atoms respectively. 500 ns long production simulations for each membrane-pore system was performed without restraints on the protein, in the NPT ensemble. System temperature was maintained at 310 K with a coupling time constant of 0.1 ps by using the stochastic rescaling thermostat. ^43^ A pressure of 1 bar was maintained by using the semi-isotropic Parrinello-Rahman barostat ^44^ (compressibilities *κ*_*xy*_=*κ*_*z*_=4.5 *×* 10^−5^ bar^-1^, time constant of 10 ps).

## 3 Results

The first part of this section is concerned with modulation of structural properties in the presence of ClyA and AHL. Properties such as the area-per-headgroup, pair correlation functions, structure factors, lipid tilt, deuterium order parameters and bilayer thickness are compared and contrasted between the two PFTs. In the last part of the structural analysis a detailed analysis of lipid protein interactions is presented. The second part of this section is concerned with the dynamics of the lipids that surround the PFT. Here in addition to the mean squared displacements of lipids we compute the displacement probabilities, continuous survival probabilities and lipid mobilities to assess the spatial perturbation in dynamical quantities surrounding the pore compex.

### 3.1 Structural Analysis

#### Area per lipid

The area per lipid (*a*_*l*_) is an important structural property strongly correlated with water permeability and transport of small molecules. ^45^ *a*_*l*_ is sensitive to lipid-lipid as well as lipid-protein interactions. The area per lipid is calculated using,

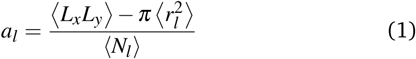

where *r*_*l*_ is the radius of the pore defined as the minimum radial distance from the center of mass co-ordinates of the corresponding PFT to the co-ordinates of the lipid phosphate head groups in the leaflet of interest and *(N*_*l*_) is the number of lipids in the corresponding leaflet. *a*_*l*_ is computed independently for both the EL and CL. Time averaged values of *a*_*l*_ evaluated for the last 350 ns of a 500 ns trajectory (see Fig. S1, ESI†) are given in Table 1. A 11% increase in the *a*_*l*_ for the ClyA bound membrane is observed in the CL while the *a*_*l*_ is relatively unchanged in the EL. The tapering structure of the ClyA pore in the CL disrupts lipid packing resulting in an increase in *a*_*l*_. However, the *β*-PFT (*α*-hemolysin) perturbs only the EL wherein one can observe a smaller 5.5% increase in the *a*_*l*_ whereas the lower leaflet is relatively unperturbed due to the *β*-barrel nature of the transmembrane segment. The cap region of AHL interacting with the EL increases the *a*_*l*_ while the hydrophobic interactions of the *β* barrel has a minimum influence on the lipid packing in the lower leaflet. Due to the different pore structures, ClyA and AHL have opposing effects on the structural rearrangement of membrane lipids. The AHL pore disrupts lipid packing in the EL while ClyA disrupts packing in the CL. We also computed the spatial variation in *a*_*l*_ by examining Voronoi diagrams constructed using the position of the lipid head groups. Interestingly, we did not observe any significant differences in the area as a function of distance from the pore complex for both ClyA and AHL indicating that the induced spatial perturbation to lipid packing is minimal.

**Table 1.** Area per lipid, *a*_*l*_ for the extracellular (EL) and cytoplasmic (CL) leaflets in the PFT bound DMPC bilayer. See text for definition of *r*_*l*_.

| System | $\langle r_l \rangle$ (nm) | $\langle a_l \rangle$ nm<sup>2</sup> |
| --- | --- | --- |
| DMPC + ClyA (EL) | 4.15 | 0.61 |
| DMPC + ClyA (CL) | 2.79 | 0.68 |
| DMPC + AHL (EL) | 1.94 | 0.64 |
| DMPC + AHL (CL) | 1.58 | 0.6 |
| DMPC (bare) | – | 0.6 |

#### In-plane Pair Correlation Function & Static Structure Factor

In order to characterize the local structure in each leaflet of the bilayer, the in-plane pair correlation function (PCF) between the head groups is calculated using,

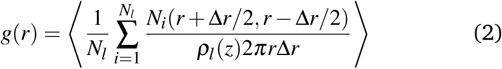

In the above equation, the areal density of the membrane is defined using the effective area, 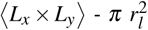 where *r*_*l*_ is the radius of pore in the corresponding leaflet of the bilayers.

The PCF of the head groups in upper and lower leaflets for ClyA and AHL are illustrated in Figs. 2a & b respectively and contrasted with the bare DMPC bilayer. An enhancement in local in-plane order is observed for the ClyA bound membrane wherein the first peak height of the EL is slightly higher than the bare bilayer. In contrast the first peak height for the CL is lower in intensity indicative of greater disorder consistent with the larger values of *a*_*l*_. Although the differences are smaller, qualitatively similar intensity differences are also observed for the second peak. Interestingly, a gradual decay in the PCF with a negative slope is observed for the lipids in the upper leaflet approaching unity around 1.5 nm, indicating the possible presence of inherent structures present in the upper leaflet possibly modulated by local packing effects. Thus the ClyA structure with the inserted hydrophobic *β* tongues ^26^ that penetrate the EL, appear to induce greater order in the surrounding lipids and the presence of the N-terminal *α* helices that make up the tapered pores in the CL has the opposite effect Fig. 2a. On the other hand, the peak heights in the PCF for both the CL and EL are very similar in the case of AHL, suggesting that in contrast to ClyA, perturbation to the lateral lipid headgroup packing is lowered in the case AHL which is a *β* PFT. Hence lateral ordering for lipids in the EL which are directly in contact with the extracellular domain of the protein do not undergo significant changes.

**Fig. 2.**
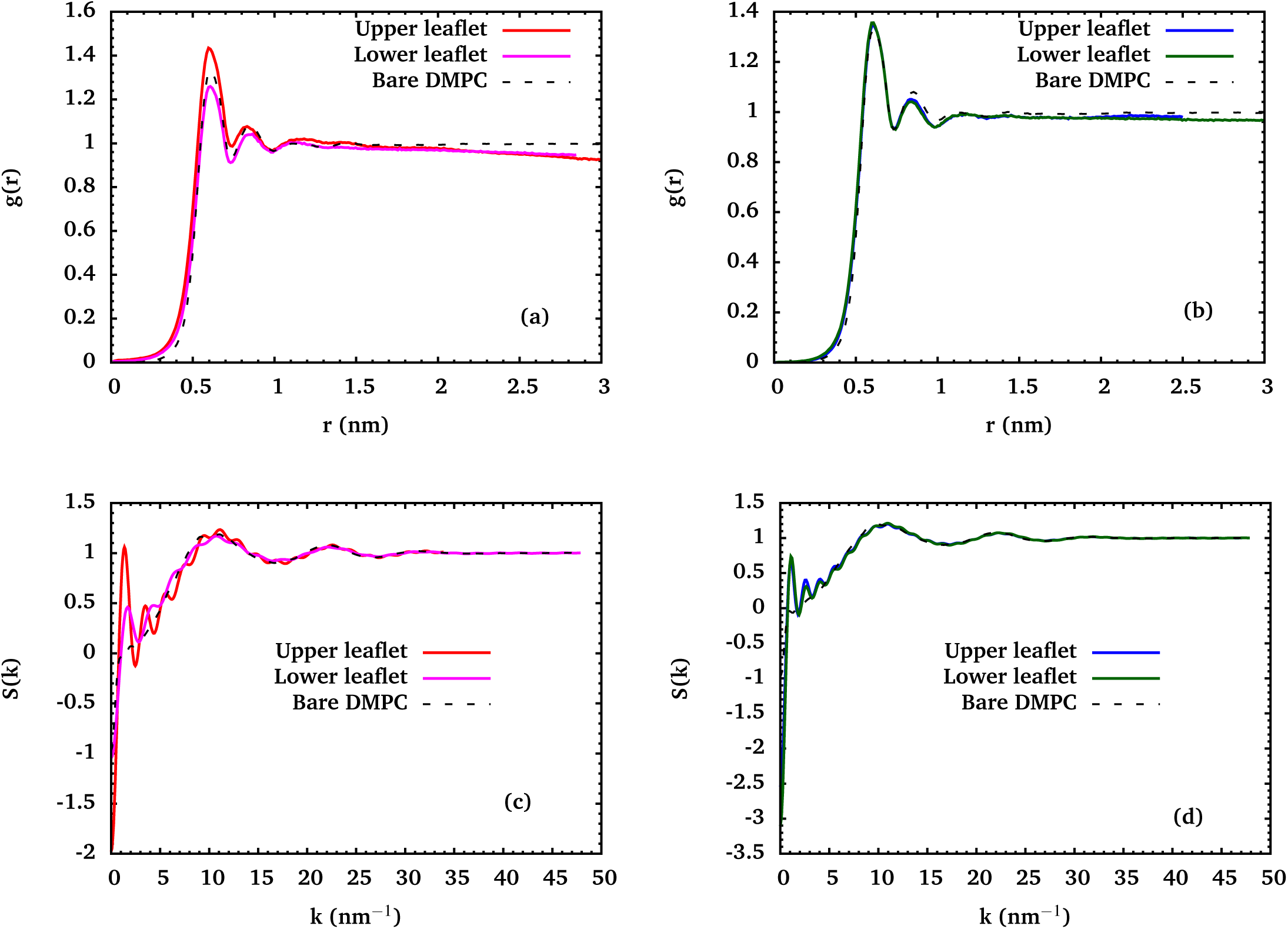
In-plane pair correlation functions between phosphate head groups of the DMPC bilayer with full pore complex formed by (a) ClyA (b) AHL. Static structure factor, *S*(*k*), evaluated between the phosphate head groups of the DMPC bilayer with (c) ClyA (d) AHL. The black dashed line represents the bare DMPC bilayer.

The corresponding static structure factors are evaluated using

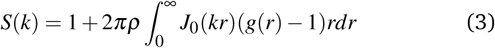

where *ρ* is the areal density of the corresponding leaflets after correcting for the pore area, *J*_0_ is the Bessel function of the zeroth order and *g*(*r*) is the in-plane pair correlation function. The Fig. 2c & d illustrate a large number of oscillations at low wavenumbers corresponding to the thermodynamic limit which are absent in the bare bilayer. The oscillations decay at larger wavenumbers. The peaks in *S*(*k*) for ClyA are sharper and closer spaced for the EL when compared with the broader peaks observed for the CL. In contrast, the period of oscillations and peak heights are similar for both the upper and lower leaflets for AHL bound bilayer. The oscillatory nature of *S*(*k*) arising from density modulations and interactions between the lipid head groups and the PFT is suggestive of a glassy state ^46–49^ of the lipid molecules induced by binding with the PFT’s. The oscillations at larger wavenumbers emerge from a finite value of the in-plane PCF at small distances. The large oscillations at small wavenumbers are indicative of long range structural correlations in the corresponding PCF’s. We note that the structure reflects the lipid in an environment consisting of an array of equally spaced pores due to the implicit periodicity present in the system (Figs. S2 and S3, ESI†).

#### Chain tilt and order parameters

The influence of PFT’s on the orientation of the lipids is investigated using the tilt angle defined for the two tails (sn1 and sn2, see Fig. S4, ESI†) of the DMPC molecule. The tilt angle is evaluated using,

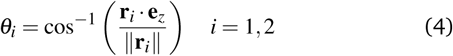

where *θ*_*i*_ is the tilt angle subtended by the vector **r**_*i*_ connecting the head (phosphorous atom) and the C-14 acyl carbon atom in each of the tails in lipid chain *i*. We also compute the order parameter ^50^ of the acyl chains,

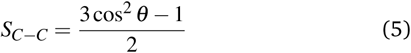

as a function of the location of carbon atoms in both lipid tails. Here *θ* is defined as the angle between the C-C bond and the bilayer normal. We investigate the acyl chain order parameter as a function of radial distance from the periphery of the protein. In the case of ClyA the inner diameter of the first shell begins at the periphery of the inserted *β* tongues (Fig. 3a) and in the case of AHL the first shell is adjacent to the inserted *β* barrel (Fig. 4a).

**Fig. 3.**
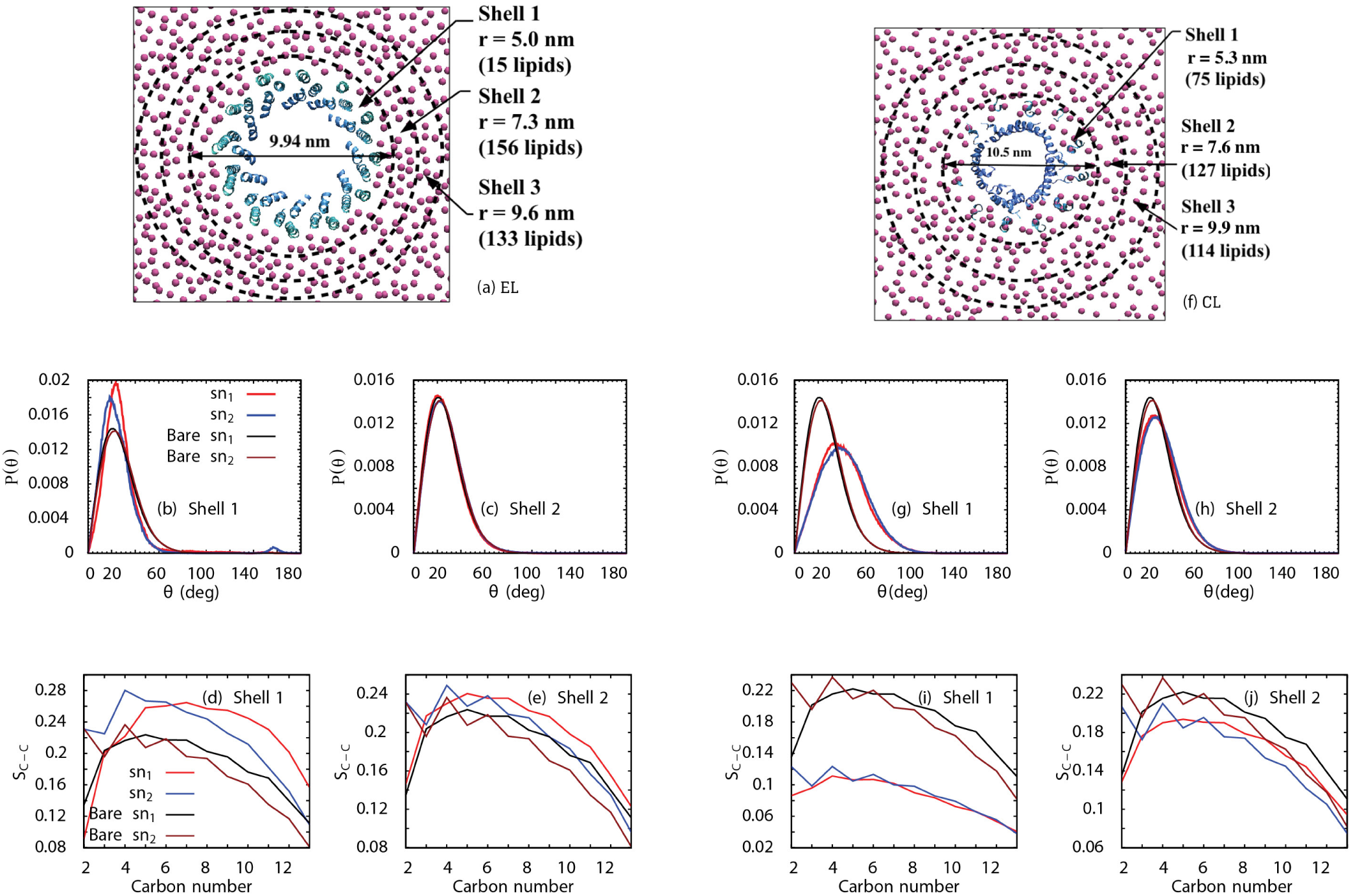
(a) Top view of the extracellular leaflet (EL) of ClyA bound DMPC bilayer divided into radial shells of thickness 2.32 nm. (b) & (c) Tilt angle distribution and the corresponding order parameter (d) & (e) in shells 1 and 2 (f) Radial shells of thickness 2.24 nm in the cytoplasmic leaflet (CL) of ClyA bound bilayer. Corresponding tilt angle distributions (g) & (h) and the order parameter (i) & (j) for the shells. Increased disruption to lipids in the EL is distinctly observed. All radial distances are measured from the center of mass of the dodecameric ClyA pore complex.

**Fig. 4.**
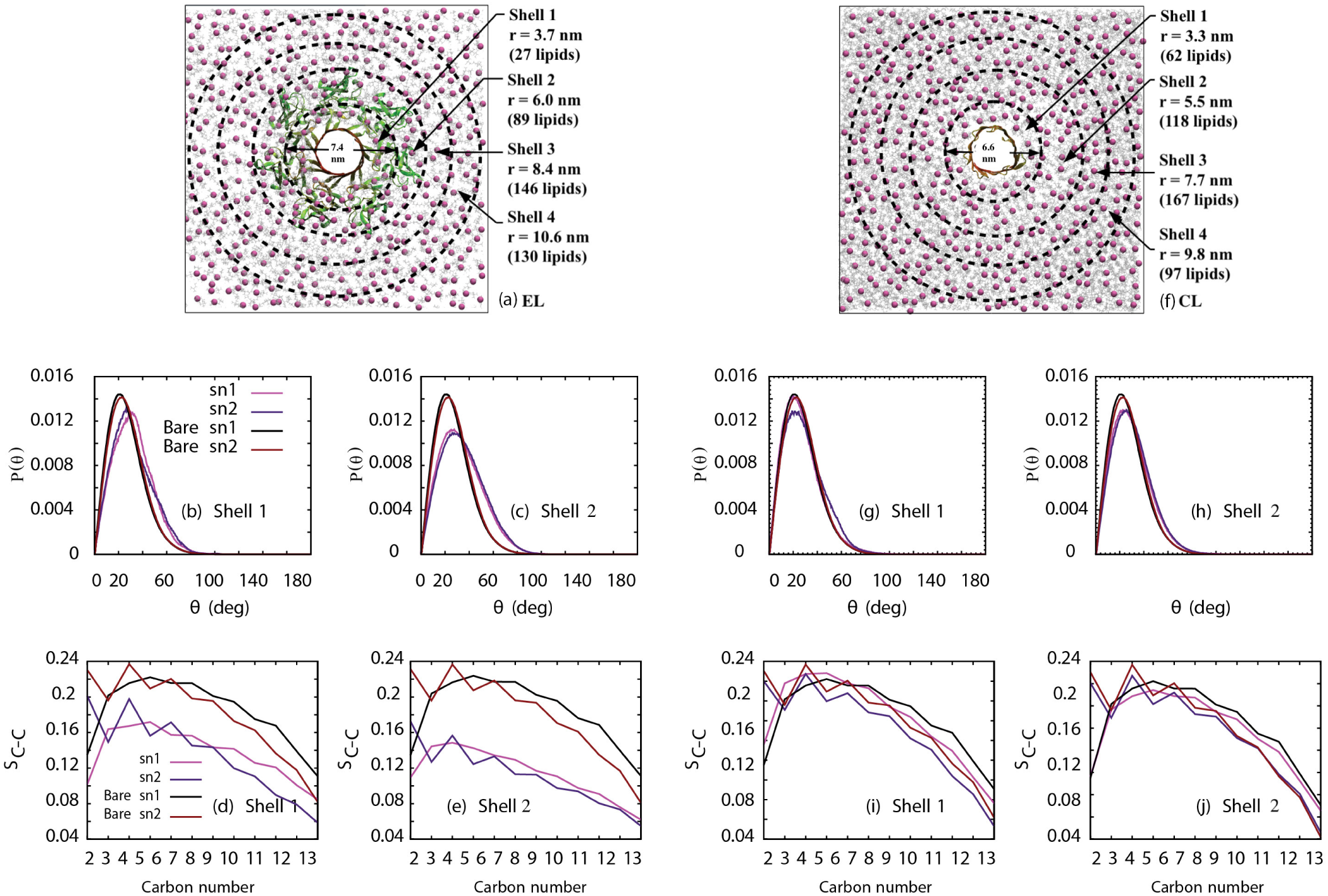
(a) Extracellular leaflet (EL) of AHL bound DMPC bilayer depicting shells of radial thickness 2.27 nm. (b) & (c) Corresponding tilt angles and (d) & (e) order parameters (f) Cytoplasmic leaflet (CL) of the AHL bound DMPC bilayer depicting radial shells of thickness 2.19 nm. Radial distances are measured from the center of mass of the pore complex. (g) & (h) Corresponding tilt angles and (i) & (j) order parameters. Ordering is disrupted in the first two shells for the EL. The tilt distribution is similar to the bare membrane in the CL, and disordering is observed in the acyl chains of the lipid tails.

The tilt angle distributions for the EL and CL of ClyA bound DMPC membrane in different shells are illustrated in Figs. 3b, c, g and h. An increase in the tilt and a narrow distribution to the bilayer normal is observed for the lipid tails in the immediate vicinity of ClyA. Interestingly the sn1 chain has a narrower distribution when compared with the sn_2_ chain, indicating a distinct asymmetry in the orientation of lipid tails in the upper leaflet due to the presence of the ClyA pore. This influence in the EL is short ranged and the tilt angle distributions rapidly approach that of the bare bilayer beginning with the second shell (Fig. 3c). In contrast to the EL, tilt angle distributions in the first two shells (Fig. 3)g & h of the CL show a significantly broadened distribution with a few lipids sampling angles greater than 100°. The recovery of the tilt angles takes places over a greater distance from the vicinity of the N-termini helices in the CL indicating that the tapered structure of the pore in the CL has a greater effect on lipid orientation.

The inserted *β* tongues in the EL predominantly consist of hydrophobic residues and hence the perturbation to the surrounding lipids is minimal. The first step in the pore forming pathway for ClyA is thought to occur with the insertion of the *β* tongue into the EL of the membrane. ^26,51,52^ During this step the *β* tongue which is in the form of a *β* sheet in the water soluble monomeric state of the protein undergoes a transformation to a membrane inserted *α* helix upon encountering the membrane. The tilt order parameter in the EL suggests that the insertion of the *β* tongue occurs with minimal disruption to the lipid tilt in the EL. Subsequent steps in the pore formation involve the insertion of the N-terminus which is an amphiphatic *α* helix to form the tapered pore structure extending into the CL of the membrane.

The corresponding *S*_*C*−*C*_ values for lipids in the EL as a function of the carbon atoms are illustrated in Fig. 3d & e. In general we observe higher values of *S*_*C*−*C*_ along the lipid chain, when compared with the bare membrane. This implies that the angle between successive C-C bonds and the bilayer normal are marginally reduced in the EL when compared with the bare membrane. This “stretching” of the lipids could be attributed to the extension of the *β* tongues below the bilayer midplane and is more pronounced for lipids in the first shell. This effect is accentuated for the sn1 tails when compared with the sn2 tails, consistent with the narrower tilt angle distribution observed for the sn1 tail. In contrast to the rapid recovery of the tilt angle distributions, the *S*_*C*−*C*_ values remain marginally higher than the bare membrane even for the shell furthest (r = 9.61 nm) from the pore center. This situation is altered for lipids in the CL. The corresponding *S*_*C*−*C*_ values for the different carbon atoms (Fig. 3i & j) illustrate the increased disorder in the lipid tails in the CL. The lowered values of *S*_*C*−*C*_ in the CL when compared with the values for the EL indicate, on average, a greater “compression” of the lipid tails in the CL and the disruption is more long ranged in the CL.

In contrast with ClyA, AHL has an extended mushroom-like cap region of the extracellular protein complex in contact with the lipid headgroups of the EL. The tilt angle distributions in the first shell of the upper leaflet (Fig. 4b & c) illustrates a shift in the peak to about 30^*°*^ for both the sn1 and sn2 tails, with broader distributions when compared with the bare bilayer. This broadened distribution for the tilt angles approach that of the bare membrane in shell 3 (not shown). Thus the overall perturbation of the chain tilt for lipids in the EL in the immediate vicinity of the AHL pore is greater when compared with the corresponding changes observed for ClyA. For AHL, only the head groups of the lipids in the upper leaflet are peripherally bound to the *β* sheet complex (cap region) which results in an extended spatial modulation of tilt angles away from the transmembrane *β* barrel pore region (Fig. 4 a). In contrast lipids in the CL have a distribution very similar to that of the bare lipid membrane even in the vicinity of the PFT (Fig. 4 (g) & (h)). There is a slight difference in the order parameter distribution between the lipid bound protein and the bare lipid membrane in the lower leaflet but a rapid recovery to bare membrane orientation occurs from shell 3 (not shown). Thus the disruption to lipid order is largely present in the upper leaflet in the case of AHL. Unlike the case of ClyA, the lipids in the CL of the AHL complex are perturbed to a lesser extent since the lipids surround the hydrophobic transmembrane *β* barrel. The *S*_*C*−*C*_ for the EL (Fig. 4d & e) indicates a lowering of the chain order resulting from a “compression” of the alkyl tails when compared with the bare membrane. This loss of ordering is observed up to the periphery of the AHL cap domain and a gradual recovery is observed thereafter. In contrast the chain order is only moderately reduced for the CL (Fig. 4i & j).

In summary, we observe that the spatial extent of disruption to the surrounding lipid structure as revealed in the tilted and *S*_*C*−*C*_ order parameters is a weak function of the distance from the pore complex. Apart from the induced asymmetry in the case of ClyA, the greatest disruption to lipid order occurs in the immediate vicinity of the pore extending to about 2.3 nm from the periphery of the pore complex.

#### Bilayer thickness

The bilayer thickness is defined as the distance between the phospholipid head groups in opposing leaflets of the lipid bilayer. If the bilayer thickness conforms with the thickness of the transmembrane PFT domain, the energy penalty involved in exposing the polar-non-polar interface is minimum. ^53^ Further, the local material properties such as bilayer compressibility, bending, splay and tilt moduli are altered with variations in thickness. The thickness maps for the ClyA and AHL bound DMPC bilayers are illustrated in Fig. 5a & b respectively and the corresponding map for the bare bilayer is given in Fig. S5, ESI†. The average bilayer thickness of the bare DMPC bilayer at 310 K is 3.46 nm. In the case of ClyA, a uniform thinning is observed in the vicinity of the pore complex extending to a radial distance of about 1 nm away from the pore periphery. Further, for the system size investigated, the bilayer thickness away from the pore was found to be lower than that of the bare bilayer indicating the spatial extent of induced height modulation with ClyA. In contrast height modulation due the AHL pore is almost absent with an induced thickness variation lying within 0.1 nm.

**Fig. 5.**
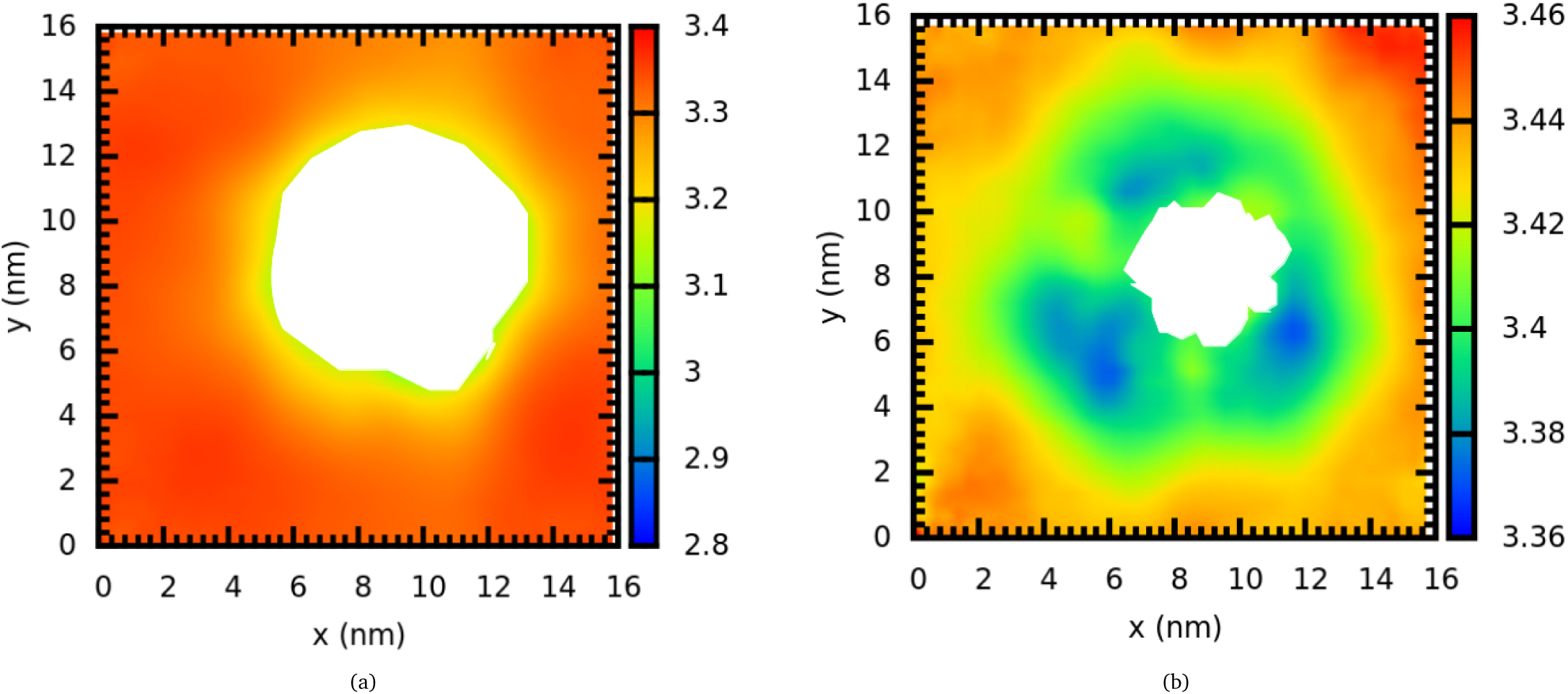
Bilayer thickness maps for (a) ClyA bound DMPC bilayer (b) AHL bound DMPC bilayer. The vertical colour bars are in units of nm. Significant bilayer thinning is observed for ClyA (a) in the vicinity of the pore complex while the membrane thickness is relatively unperturbed for AHL (b). The white region depicts the pore complex.

The larger hydrophobic mismatch in the case of ClyA in DMPC implies a greater energetic penalty during the binding of this *α*-helical protein with its wedge shaped membrane inserted topology. This is evidenced by the extent of membrane thinning observed for the ClyA membrane in the vicinity of the pore complex. In order to minimize the energetic and mechanical penalty associated with this hydrophobic mismatch, a substantial broadening in the tilt angles occurs for the CL (Fig. 3) and the *S*_*C*−*C*_ also reveal that lipid chains in the EL (Fig. 3) are “stretched” while lipids in the lower leaflet are “compressed” relative to the bare DMPC bilayer. In contrast the hydrophobic mismatch for AHL in the DMPC membrane is virtually absent, indicating the reduced energetic penalty for pore formation for this *β* PFT.

#### Lipid-protein interactions

The structural properties analysed in previous sections clearly reveal the inherent asymmetry induced in the EL and CL of the DMPC bilayer by the presence of the different PFTs modulated both by the chemical and geometric nature of the inserted secondary structure for ClyA and AHL. In order to shed insights into the observed leaflet induced asymmetry as observed in several of the structural quantities such as the pair distribution functions, tilt and deuterium order parameters and thickness maps, we evaluate the difference in the number of atomic contacts, contributions from non-polar and polar contacts, as well as the hydrogen bonding of specific protein residues on the interactions with the lipids in the EL and CL. The number of unique atomic contacts between two non-overlapping groups, such as between the protein and membrane lipids, was calculated by using the Gromacs tool ‘mindist’ with a default distance criterion of 0.6 nm.

The number of atomic contacts between the protein and lipid for both the CL and EL as well as their contributions from non-polar and polar contacts are illustrated in Fig. 6 for both ClyA and AHL. In panels c, d, e and h, the largest number of atomic contacts is scaled to 100. ClyA shows a 12% higher number of contacts with the CL while AHL strikingly shows a 65% higher number of contacts with the EL. The nature of protein-lipid contacts are also distinctly different between the membrane leaflets (Fig. 6e & f)). Both ClyA and AHL pores predominantly engage in non-polar residue-lipid contacts with the CL (79.4% and 84.4% for ClyA and AHL, respectively), whereas the number of polar and non-polar residue-lipid contacts are approximately equal in the EL. Region-wise breakdown of the ClyA and AHL membrane contacts into headgroup, sn1 acyl chain, and sn2 acyl chain contacts in the CL and EL highlight the difference in the nature of ClyA and AHL membrane interactions (Fig. 6g & h). The higher number of contacts with the CL by ClyA is mostly due to non-polar residue-lipid contacts with the sn1 and sn2 tails. In contrast, the higher number of contacts with the EL by AHL is due to a steep increase in the number of protein-headgroup contacts.

**Fig. 6.**
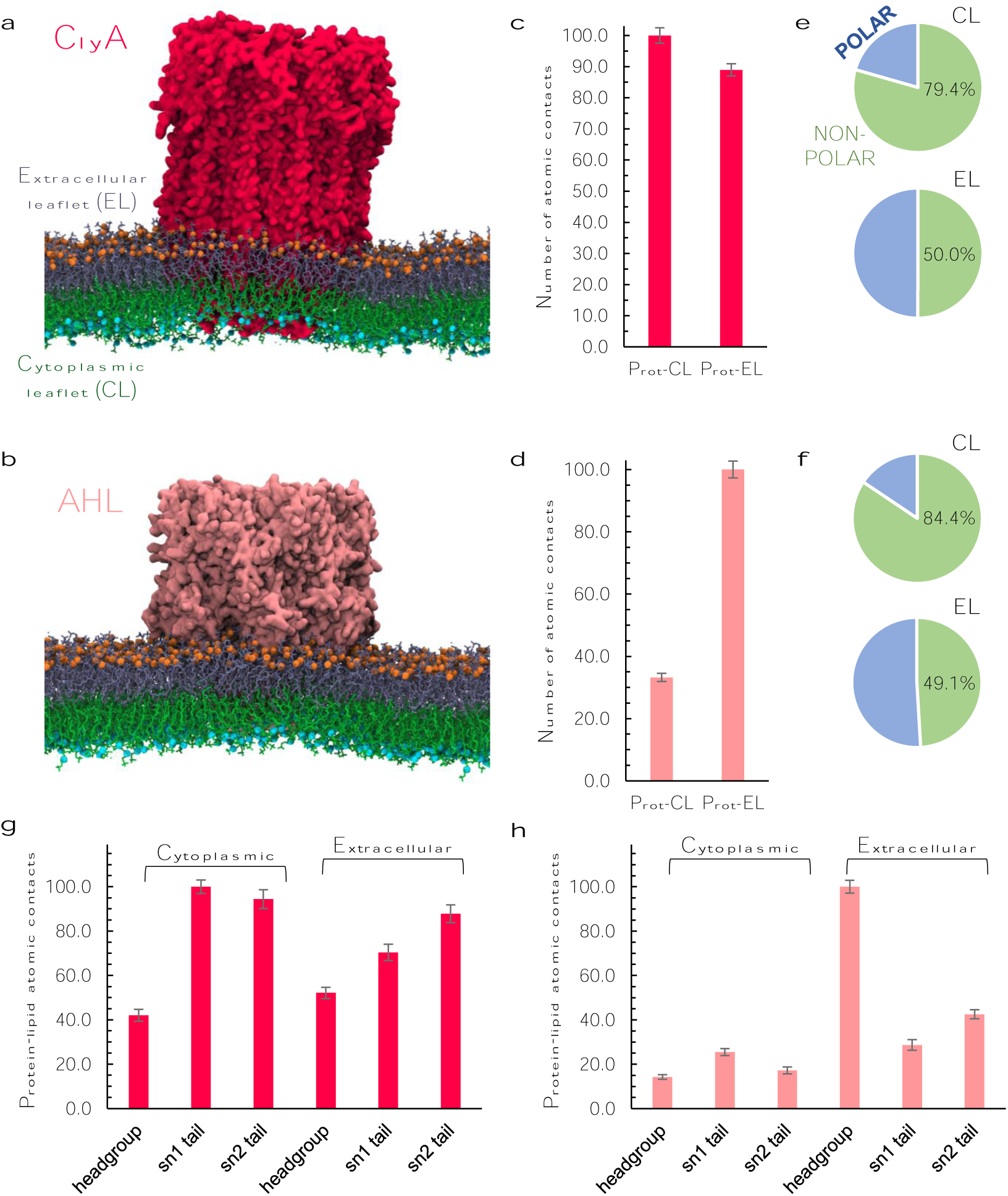
Simulation snapshots for the transmembrane ClyA (**a**) and AHL (**b**) pores at 500 ns; ClyA and AHL pores coloured red and pink, respectively, extracellular leaflet (EL) coloured violet, and cytoplasmic leaflet (CL) coloured green. Phosphate beads in the extracellular and cytoplasmic leaflets are coloured orange and cyan, respectively. **c, d** Number of contacts between the ClyA (**c**) and AHL (**d**) protein (Prot) atoms and EL/CL atoms averaged over the last 50 ns of the MD trajectory. **e, f** Fraction of protein-lipid contacts by polar and non-polar amino acids in the extracellular and cytoplasmic leaflets by ClyA (**e**) and AHL (**f**) pores, respectively. **g, h** Region-wise breakdown of the protein-membrane atomic contacts into lipid headgroup-protein, sn1 hydrocarbon tail-protein and sn2 hydrocarbon tail-protein contacts, respectively, for the ClyA (**g**) and AHL (**h**) pores. In all cases the contacts are normalized to 100 using the largest number of contacts for each case. The actual number of contacts are given in Table S1, ESI†.

In the case of ClyA, protein-lipid contacts in the EL arise predominantly between the inserted helices formed by the *β* tongues in the EL of the membrane. The inserted *β* tongues consist predominantly of non-polar residues (green) as illustrated in Fig. 7a & c, which interact with lipid tails giving rise to the non-polar protein-lipid contacts in the EL. In contrast the amphiphatic helical bundle made up of N-terminal *α* helices present in the CL, has a larger number of non-polar lipid-protein contacts. The differences in the geometry of the two pore architectures influences the partitioning of the atomic contacts. As a consequence protein-lipid contacts dominate in the case of ClyA (Fig. 6g) for both leaflets due to the inserted transmembrane helices. However, in the case of AHL, the extended cap domain in addition to the *β* barrel contribute to the increased number of contacts in the EL, when compared with the CL where only *β* barrel-lipid contacts are present. Hence enhanced protein-lipid headgroup contacts dominate over the protein-lipid tail contacts in the EL (Fig. 6h).

**Fig. 7.**
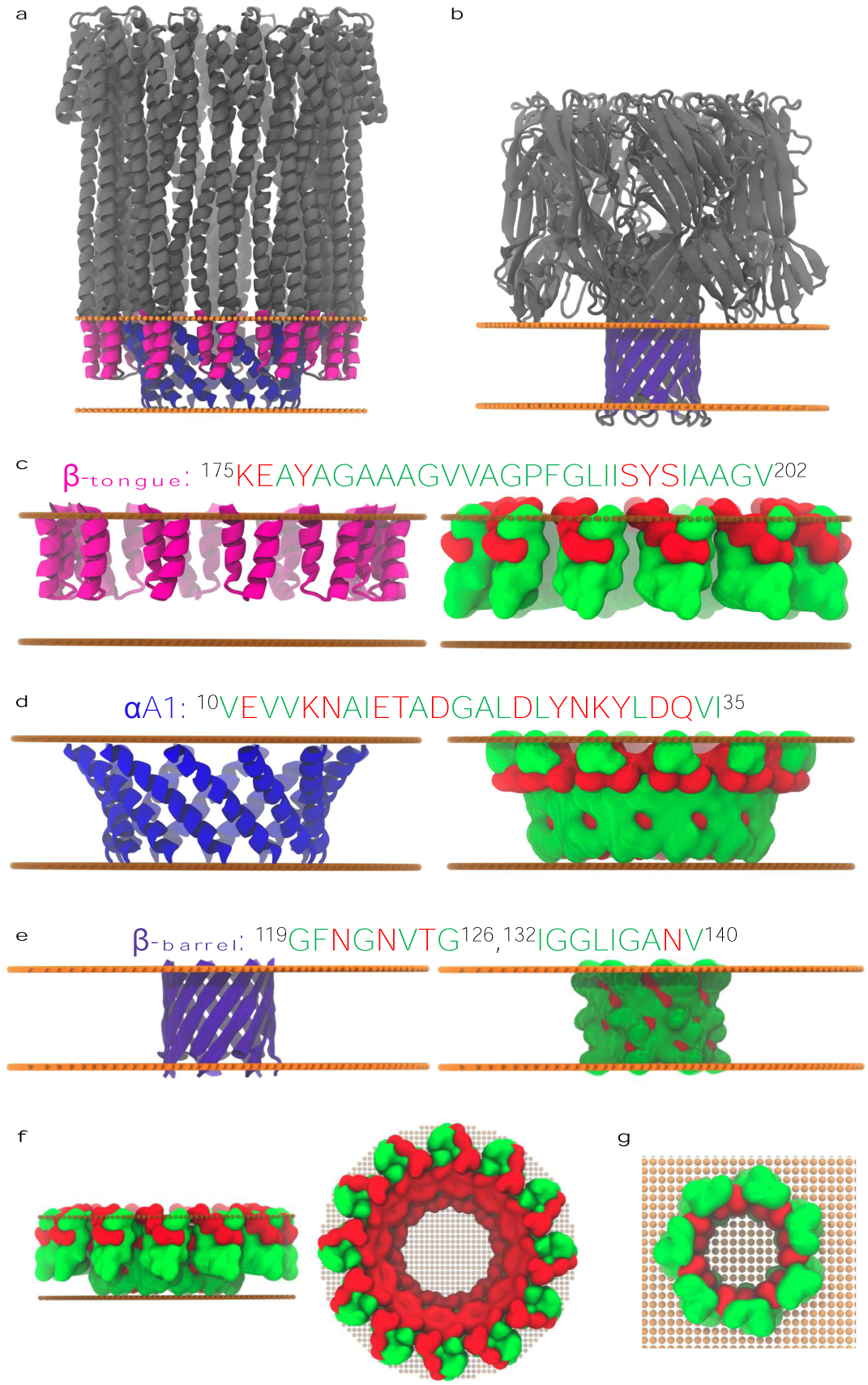
Transmembrane domains of (**a**) ClyA (*α*-helix A1, coloured blue, residues 10-35; *β*-tongue, coloured magenta, residues 175 to 202) and AHL (*β*-barrel, coloured violet, residues 119-126, 132-140) are illustrated (solvated residues in both pores are coloured grey). Amino acid sequence along with cartoon and corresponding surface representations of (**c**) the *β*-tongue in ClyA (**d**) the *α*-helix A1 in ClyA and (**e**) the *β*-barrel in AHL are shown. Green and red colour in the sequences and the surface representations indicate polar and non-polar residues, respectively. (**f**) Front and top views of the entire transmembrane domain of ClyA (*α*-helix A1 + *β*-tongue) are shown with surface representation. All of the lipid interacting residues are non-polar (except possibly at the extracellular membrane-water interface) while the inner pore lumen is almost completely hydrophilic. (**g**) Top view of AHL. Similar to ClyA, the outer membrane interacting surface is mostly non-polar while the inner pore lumen is hydrophilic.

Since protein residues that lie on the periphery of the membrane-water interface play a key role in anchoring the PFT to the membrane, we evaluate the interaction of a few amino acid residues that have been shown to modulate interactions of the protein with the headgroups and acyl chains of the membrane lipids. ^23,24^ Two classes of amino acids in membrane proteins effect the structure of the surrounding membrane through interactions with the lipid head groups: the amphipathic aromatic amino acids, tyrosine (Y) and tryptophan (W), and the basic amino acids, lysine (K) and arginine (R). Tyrosine and tryptophan anchor and orient membrane proteins in the bilayer through hydrogen bonding and cation-*π* interactions with the lipid head groups as well as favorable amino acid-dipole interactions with the strong membrane-water interfacial electric field. ^54–56^ The basic amino acids, lysine and arginine, facilitate the membrane penetrative function of membrane disruptive proteins ^57,58^ and stabilize the structure of the membrane-protein-water interface. Arginine with its five potential hydrogen bond donors and high pKa can form a variety of complex hydrogen bonding patterns such as the arginine-fork ^59^ and the arginine guanidinium-lipid phosphate complex ^60^ with the surrounding lipids. Similarly, lysine is important for membrane binding ^61^ and is shown to have a specific affinity to the lipid phosphate groups near the aqueous interface. ^62^ Arginine and lysine extensively hydrogen bond both with water and the polar lipid phosphate and glycerol groups. Additionally, arginine and lysine engage in ‘snorkelling’, where the aliphatic parts of the side chain prefer localization in the membrane core due to hydrophobic interactions and the positively charged side chain terminus interacts enthalpically with the membrane-water interface. ^63^ The exact location of these membrane-interacting ‘amino acid belts’ therefore significantly influences the orientation of membrane proteins as well as the surrounding membrane structure through interfacial residue interactions with specific lipid moieties.

The RKWY contacts help to orient and anchor into the membrane and therefore, we calculated possible RKWY protein-lipid interactions in the transmembrane ClyA and AHL pores (Fig. 8). Remarkably, both ClyA and AHL pores show ∼ 3.2-fold and ∼ 24.1-fold higher RKWY contacts with the extracellular leaflet compared to the cytoplasmic leaflet, respectively (Fig. 8a). Both ClyA and AHL predominantly form protein-lipid hydrogen bonds with the extracellular leaflet (Fig. 8b). Of these, protein-membrane hydrogen bonds by RKWY residues account for ∼ 100% in ClyA and ∼ 53% in AHL, respectively (see Fig. 8b). The major hydrogen bonding RKWY residues, R174, Y178 and K206 in ClyA are located in the vicinity of the *β*-tongue region, which is in close proximity to the lipid-water interface, and their hydrogen bond interactions with the lipids in the 500 ns MD snapshot are illustrated in Fig. 8c. These residues have also been implicated in strong interactions with cholesterol. ^51^ Similarly, the RKWY residues Y118, W187, R200 and K266 form the bulk of the interactions in the AHL pore. These RKWY amino acid belts stabilize the extracellular leaflet in both ClyA and AHL pores, and may significantly influence structural and dynamical asymmetry across membrane leaflets. In addition to hydrogen bonds with the extracellular leaflet, a substantial fraction of RKWY-EL contacts are also with the sn1 and sn2 tails, which is indicative of snorkelling and hydrophobic interactions. These interactions are substantially lower in the cytoplasmic leaflet.

**Fig. 8.**
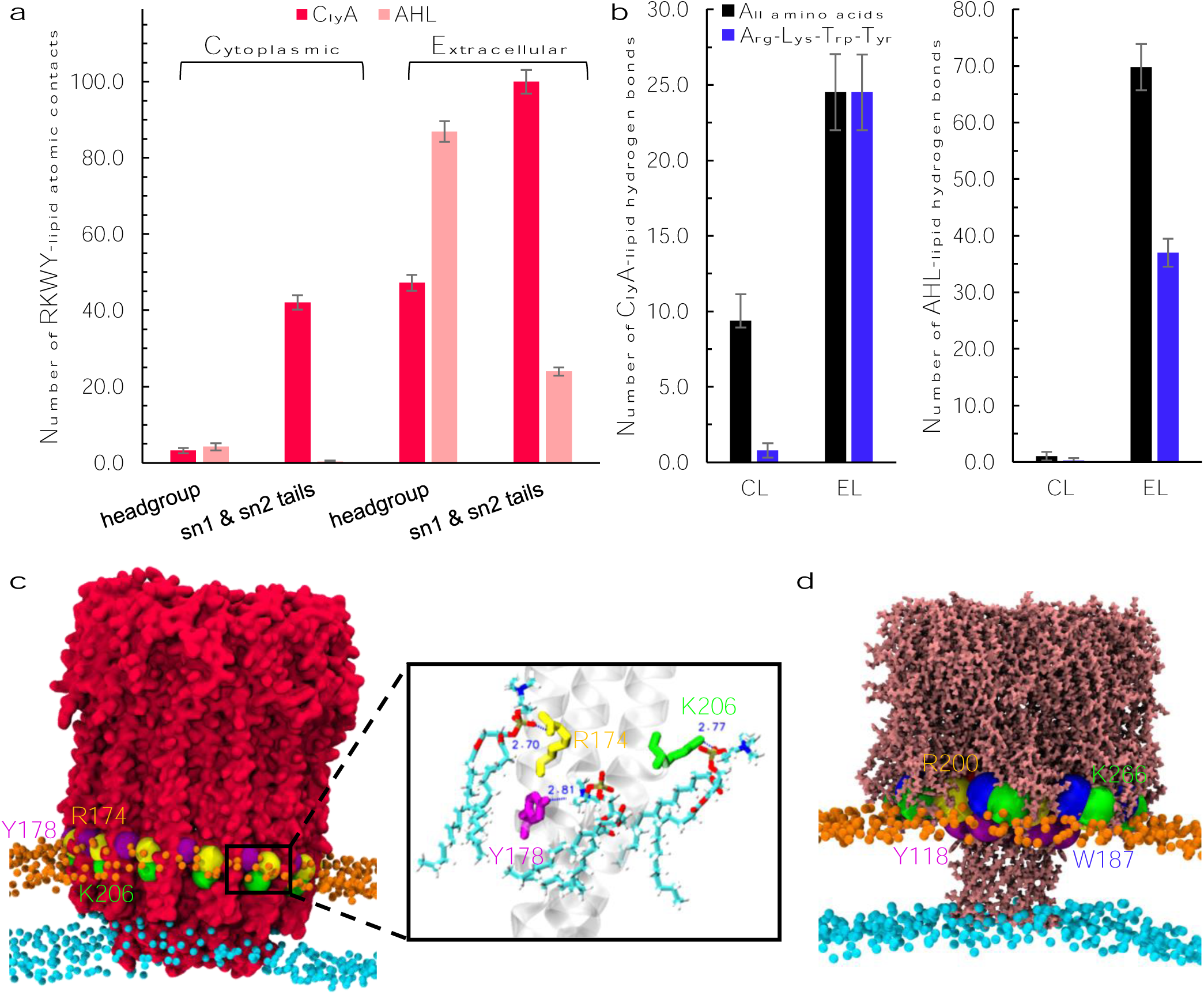
**a**, Region-wise breakdown of EL/CL atomic contacts into lipid headgroup and hydrocarbon tail contacts with the membrane-interacting amino acids arginine (R), lysine (K), tryptophan (W) and tyrosine (Y) for the ClyA and AHL pores. **b**, Number of protein-membrane hydrogen bonds by all the ClyA and AHL protein residues (black) or just the RKWY residues (blue), separately with EL and CL. The numbers are averages over the last 50 ns of the MD trajectory. **c**, Residues R174 (yellow), Y178 (purple) and K206 (green) in the ClyA pore form strong hydrogen bonds with the extracellular leaflet lipids in the interfacial region and are depicted as spheres. Individual residue-lipid hydrogen bonds are illustrated in the magnified image on the right. **d**, Residues Y118 (purple), W187 (blue), R200 (yellow) and K266 (green) in the AHL pore form interact strongly with the extracellular leaflet lipids via hydrogen bonding in the interfacial region, and are illustrated as spheres. In **a** the contacts are normalized to 100 using the largest number of contacts. The actual number of contacts are given in Table S1, ESI†.

### 3.2 Dynamical Properties

#### Mean squared displacements

The mean squared displacement, MSD of the lipids is calculated using,

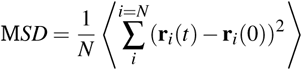

where **r** = *r*_*x*_**e**_*x*_ + *r*_*y*_**e**_*y*_. The position vector **r**_*i*_ of the lipid molecule ‘*i*’ represents positions after removing the center-of-mass motion of the individual leaflets. The EL and CL leaflets are distinguished from the mid-plane as observed in the density distributions and the corresponding center-of-mass co-ordinates are computed considering both the protein and lipid molecules in each of the leaflets.

The MSD data of the ClyA and AHL membranes are illustrated in Fig. 9a & b respectively. Lipid dynamics is marginally retarded for the upper leaflet of the ClyA membrane when compared with the lower leaflet. Since the MSD ∼ *t*^*a*^, the power law exponents, *a* are represented as *α, β* and *γ* to illustrate different dynamical regimes. These regimes are illustrated in the data for each of the leaflets in Table 2 where different time windows obtained from slopes of the MSD versus time data are compared with data obtained for the bare bilayer. The values of exponents *a* are similar to those reported in earlier MD simulations of lipid bilayers. ^64,65^ In the case of ClyA, an extended sub-diffusive regime is observed for the CL, whereas over the 300 ns sampling window the exponent approaches unity for the EL. This suggests that although the MSD is greater in the CL due to the increased disorder arising from the tapering N-termini helices, the system dynamics has greater heterogeneity as revealed from the smaller values of the exponent. The opposite is true for lipids in the EL where the diffusive regime is observed for *t >* 40 ns. The lowered dynamical heterogeneity (deduced from the exponents) in the EL indicates decreased perturbation due to the membrane inserted *β*-tongue helices. Extended simulations with larger sample sizes are required to reliably extract a diffusion coefficient and we did not pursue this aspect here.

**Table 2.** Exponents of the various sub-diffusive regimes in the MSD data of the extracellular leaflet, EL and cytoplasmic leaflet, CL. MSD ∼ *t*^*a*^, where *a* ≡ *α, β* and *γ* to differentiate between the different regimes. For the bare bilayer in the absence of protein, *α* = 0.548 (0.004 ≤ *t* ≤ 4 ns), *β* = 0.858 (4 ≤ *t* ≤ 220 ns) and *γ* = 1 (220 ≤ *t* ≤ 350 ns).

| Time (ns) | EL (DMPC + ClyA) | Time (ns) | CL (DMPC + ClyA) |
| --- | --- | --- | --- |
| 0.01 - 6 | $\alpha = 0.5665 \pm 0.0017$ | 0.01 - 6 | $\alpha = 0.5439 \pm 0.0034$ |
| 6 - 40 | $\beta = 0.8622 \pm 0.0033$ | 6-300 | $\beta = 0.841 \pm 0.001$ |
| 40 - 300 | $\gamma = 1$ | — | — |
| Time (ns) | EL (DMPC + AHL) | Time (ns) | CL (DMPC + AHL) |
| 0.01 - 7 | $\alpha = 0.562 \pm 0.003$ | 0.01 - 8 | $\alpha = 0.584 \pm 0.0035$ |
| 7 - 300 | $\beta = 1$ | 8 - 320 | $\beta = 1$ |

**Fig. 9.**
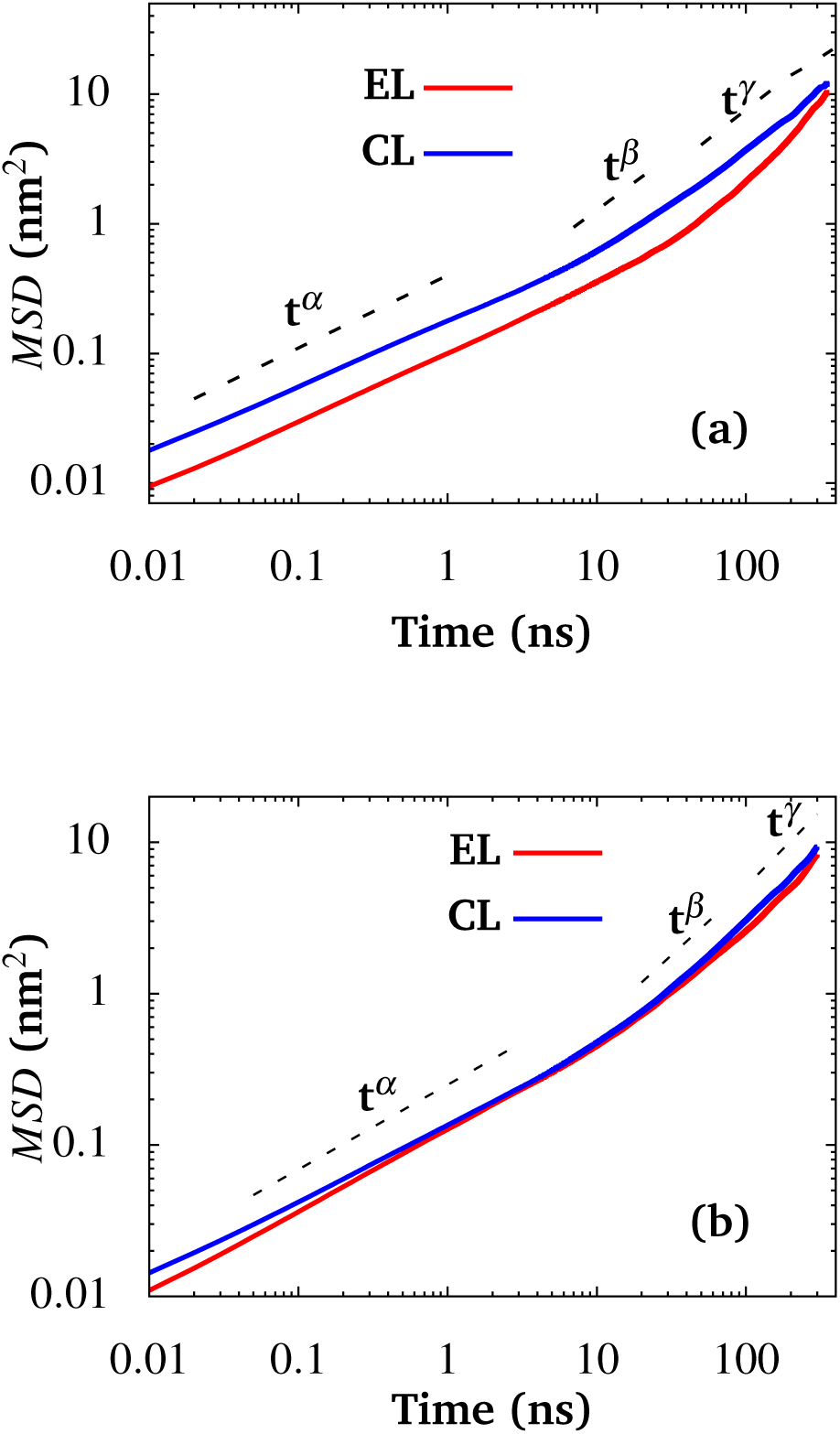
Mean squared displacement (MSD) of DMPC bilayer with (a) ClyA (b) AHL illustrating leaflet specific dynamics.

For AHL (Fig. 9b), MSD data for lipids in both leaflets are similar achieving an exponent of unity for t > 8 ns. The presence of the extended cap for AHL does not significantly perturb the dynamics in the EL when compared with the dynamics in the CL. Furthermore, although the membrane inserted *β* barrel modulates the lipid tilt and *S*_*C*−*C*_ (Fig. 4), the induced dynamic heterogeneity across leaflets is minimal.

Upon comparing the exponents for the bare bilayer, the duration of the sub-diffusive regimes in the presence of PFTs appear to be less extended suggestive of reduced dynamic heterogeneities in the presence of the membrane inserted protein complex. The induced heterogeneity and extent of the sub-diffusive regime is lower for AHL when compared with ClyA in the timeframe of the simulations explored.

#### Continuous Survival Probability (CSP)

The continuous survival probability (CSP) of the lipids was calculated in the different radial shells to understand the residence time of lipids. The construction of these radial shells is similar to those used for the order parameter calculations. At any time *t*, the CSP of *N*_*j*_ lipids in radial shell *j* is evaluated using,

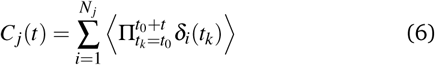

where *δ* is the Kronecker delta which is either unity if the coordinates of the phosphate head group is present in the corresponding shell or zero if absent. The CSP at any time instant ‘*t*’ is *C*_*j*_ (*t*)*/C*_*j*_ (*t*_0_).

For ClyA a distinct two-step relaxation is observed (Fig. 10a) for lipids in the EL: (i) An early relaxation wherein the decay time decreases with increasing distance from the periphery of PFT and (ii) a plateau regime extending from 10 −300 ns. In the case of CL (Fig. 10b), the early relaxation is followed by a weak plateau regime which rapidly decays. The relaxation time of both the early and late plateau regimes decreases with increasing distance from the periphery of ClyA. A key point to note is that, over the course of 300 ns majority of the lipids are present in their original locations indicative of high residence times and retarded dynamics due to the transmembrane pore complex. In the EL only 93-96% of lipids remain in shells (1-2) which are in close proximity to the membrane inserted *β*-sheets and even for the ultimate shell (shell 3) about 91% of lipids remain in this region indicating the influence of strong protein-lipid interactions. However, a more rapid displacement of lipids occurs in the CL with about 75-80% of lipids remaining at 300 ns. These trends are consistent with the increased disorder associated with lipids present in the CL as revealed in both the tilt order parameter and lowered *S*_*C*−*C*_ (Fig. 3i & j) as well as with the increased MSD observed for lipids in the CL (Fig. 9a).

**Fig. 10.**
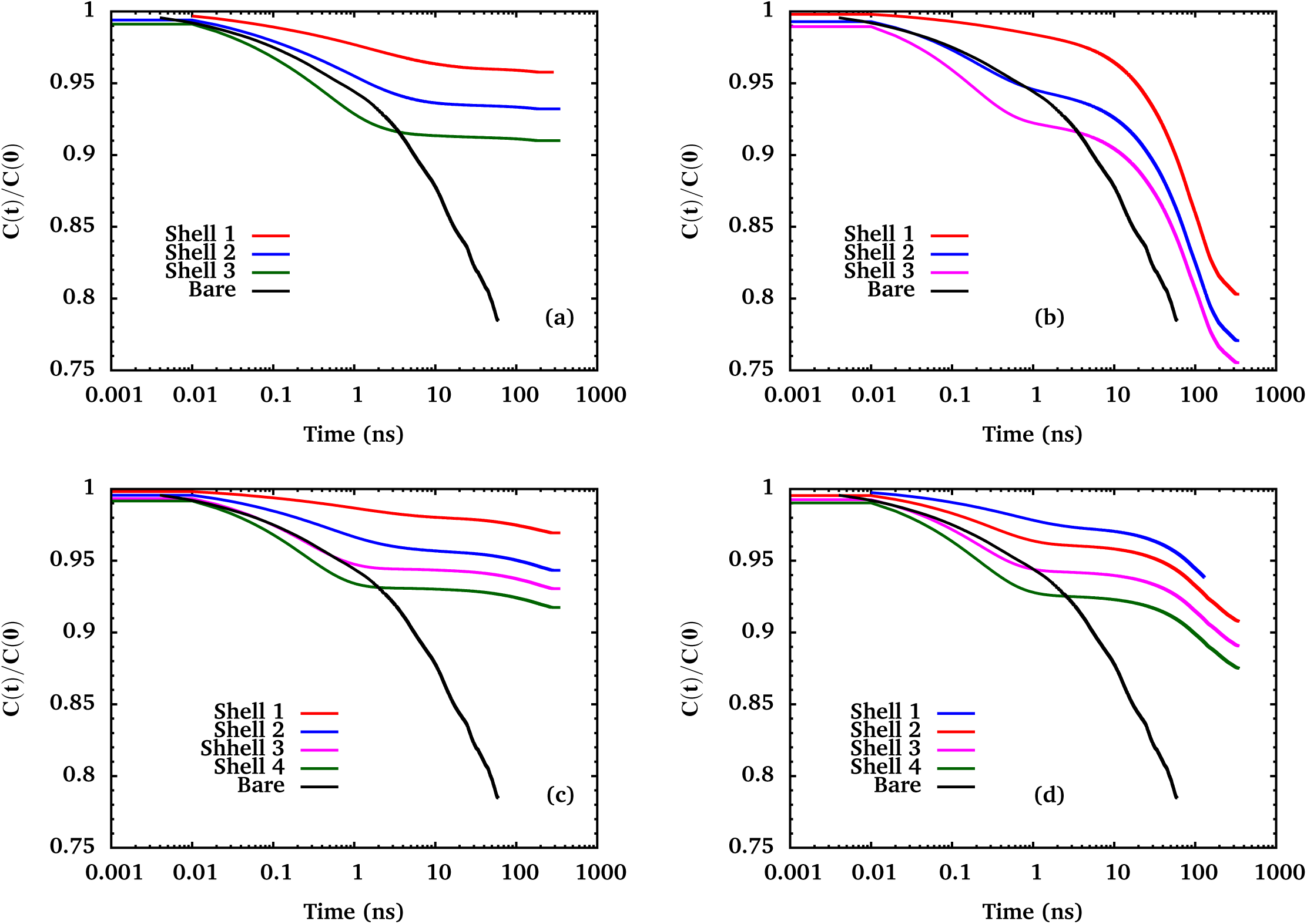
Continuous survival probability of ClyA bound DMPC. (a) Extracellular Leaflet (b) Cytoplasmic Leaflet. Continuous survival probability of *α*-hemolysin bound DMPC (c) Extracellular Leaflet (d) Cytoplasmic Leaflet. The layers are progressively numbered from the periphery of PFT.

In the presence of AHL, the lipid dynamics is arrested to a greater extent when compared with the ClyA membrane. Almost 97% of the lipid molecules continue to reside in the first shell and similar relaxation trends reveal that lipids in contact with the extracellular cap domain of AHL have a higher residence time when compared with lipids away from this region. Thus a distinct signature of the slow dynamics of lipids in contact with the AHL cap is observed. In contrast, the CSP begins to display a two step relaxation from second shell onwards, however about 91-94% of the lipids continue to reside in this region. For the CL (Fig. 10d) lipids in the first shell shows a weak two step relaxation which gets more pronounced further away from the protein, revealing the influence of the *β* barrel on the residence time dynamics and only about 10% of the lipids exit in this leaflet. With the exception of lipids in the CL of the AHL membrane, a common feature is the high residence times observed for the lipids, with only about 10% of lipids found to leave the vicinity of the proteins over the course of 300 ns. Thus lipids even the last shell which is about 10 nm from the pore center of mass or about 4 nm from the surface of the pore, possess retarded translational dynamics indicating the expanded region of dynamic slowdown due to the pore complex. The increased residence time in the EL is consistent with the larger number of protein (RKWY)-lipid contacts and greater protein-lipid hydrogen bonds in the EL when compared with the CL as illustrated in Fig. 6 & Fig. 8). Thus stronger electrostatic interactions coupled with lipid-protein interactions influence the residence times of lipids in these systems, leading to a dynamically heterogeneous environment. These results indicate the presence of a tightly bound shell of lipids around the protein complex and several *mu* s long simulations would be required to elicit a reliable relaxation time from the survival probabilities.

#### Displacement Maps and Probability Distributions

In order to assess the extent of dynamic heterogeneity as revealed in the CSP data, we evaluate the displacement maps in this section. The displacement, *µ*_*i*_ for lipid ‘*i*’ based on the displacement over a time interval Δ*t* is,

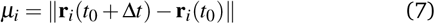

where a value of Δ t = 50 ns was used and **r**_*i*_ is the position of the phosphate head group of the corresponding lipid. Fig. 11a & b illustrates the displacement maps for lipids in the CL and EL for the ClyA bound membrane. We note that although the range of mobilities is about 1 nm for Δ*t* = 50 ns the extent of spatial heterogeneity is clearly discerned. At this time scale the exponents in the MSD data (Table 2) are greater than 0.8 for all the systems investigated. In both the CL and EL a distinct circular region of lowered displacements extends over a radial distance of about 1.5 nm from the periphery of the pore. In general, mobilities are higher for the CL, consistent with the lower residence times revealed in the corresponding CSP data for ClyA in the CL (Fig. 10b). Similar trends are observed in the displacement maps for the AHL bound membrane (Fig. 11e & f) with reduced displacement in the vicinity of the pore. For the EL, the region of lowered displacement corresponds closely to the radial extent of the cap domain (≈ 2 nm) from the *β* barrel. Interestingly the range of mobilities is quite similar for both EL as well as CL of AHL. Similar slowing down of cholesterol and lipids in the vicinity of the ClyA pore complex in POPC membranes has recently been observed by us. ^51^

**Fig. 11.**
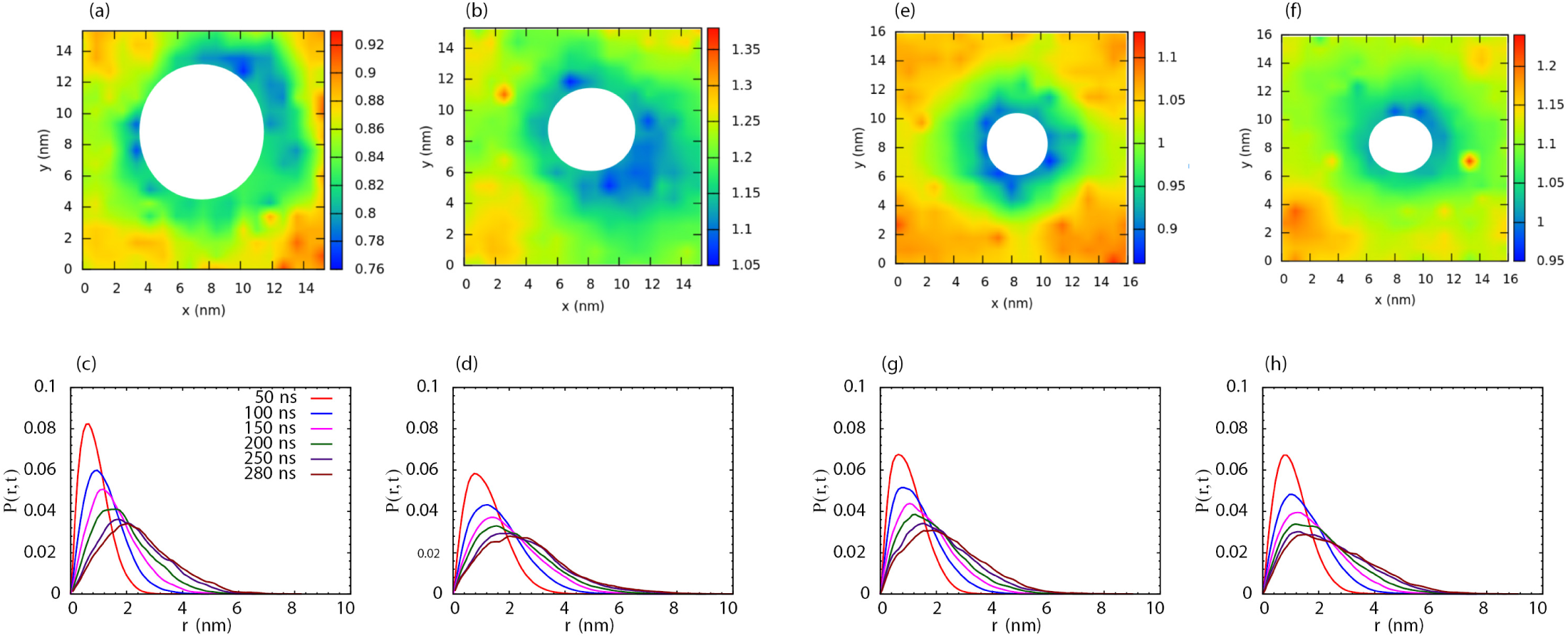
Displacement maps (top row) and displacement probability distributions, *P*(*r, t*) (bottom row). (a) and (c) EL for ClyA, (b) and (d) CL for ClyA. (e) and (g) EL for AHL (f) and (h) CL for AHL. Distinct non-Gaussian nature of *P*(*r, t*) is observed in all cases indicative of the inherent dynamic heterogeneity in the lipid dynamics. Color bars are in units of nm.

Fig. 11c & d illustrates the probability distribution of lipid displacements for the EL and CL of the ClyA inserted membrane.The sampling of phase space is greater for the CL, characterized by lower peak heights and broader distributions (Fig. 11b) at all times reflecting larger displacements in the displacement maps. The reduced overlap and broadening (non-Gaussian behaviour) of the distribution at large *r* illustrates the presence of more than one sub-diffusive regime, a signature of anomalous diffusion ^66^, as observed in the corresponding MSD data (Fig. 9a & b). Distinct non-Gaussian nature of *P*(*r, t*) for t *>* 200 ns suggest that lipid molecules are trapped in cages unable to fully sample phase leading to dynamic heterogeneities and glass-like behaviour. ^66,67^ In this regime, relaxation is predominantly governed by co-operative motion of cages wherein lipid molecules are trapped by their neighbours. A similar observation of the sampling of phase space with time can be drawn for AHL bound DMPC bilayer as well (Fig. 11g & h), however, the peak heights and spread of the curve are comparable for both the leaflets at all times unlike the ClyA bound bilayer wherein, distinct asymmetry is observed in displacements of the individual leaflets. This observation is consistent with the corresponding MSD data (Fig. 9b). Nonetheless, the system has not sampled the diffusive regime and displacement probabilities for the entire bilayer are qualitatively similar to the non-Gaussian behaviour observed for the ClyA membrane. We further point out that the slope of the MSD curves becoming linear with time is necessary but not a sufficient condition to extract a self-diffusion coefficient as observed from the *P*(*r, t*) trends for these systems. The non-Gaussian nature of the distributions for both ClyA and AHL indicate that the dynamics in these protein-lipid complexes are heterogeneous similar to glass-like systems where caging and pinning effects ^68^ are likely to play a role.

## 4 Discussion

Pore forming toxins, widely implicated in bacterial pathogenesis, mount their virulence by secreting proteins that bind to target mammalian cells forming transmembrane pores that eventually lyse the cells. In this manuscript we investigate the influence of two main classes of PFTs and their influence on the properties of a phospholipid bilayer. All atom molecular dynamics simulations are carried out for Cytolysin-A, ClyA which is an example of an *α* PFT and for *α*-hemolysin, AHL which is an example of a *β* PFT in a DMPC membrane. These two PFTs whose crystal structures have been reported, confer distinct proteo-lipidic interactions and we systematically investigate the variation in structural and dynamical properties of the lipids to assess the extent of spatial modulation induced by these proteins.

### 4.1 Leaflet Induced Asymmetry

In the case of the dodecameric ClyA pore complex, the inserted helices of the *β* tongue sample the EL forming a circular ring which define the boundaries of the pore complex. Within this ring lies the helical bundle formed by the alpha helices of the N-termini which form a truncated cone tapering into the CL (Fig. 7). An analysis of the lipid-protein contacts reveal about 50% greater polar contacts in the EL when compared with the CL. Enhanced contact with RKWY residues in the protein and lipids are observed for the EL with increased contacts with alkyl tails when compared with the head groups. Thus the penetrated transmembrane helices in ClyA provide a conducive environment for enhancing protein-lipid and hydrogen bond interactions with the EL in comparison with the CL. As a consequence lipids in the EL have enhanced order as revealed in the *g*(*r*), chain tilt and *S*_*C*−*C*_, when compared with lipids in the CL where lipids sample a broader range of tilt angles with increased lipid chain disorder. This also results in enhanced dynamic heterogeneity in the CL when compared with the EL as reflected in the MSD and *P*(*r, t*) Perhaps the single most discerning characteristic between AHL and ClyA is the reduction in the extent of induced structural and dynamic heterogeneity between the EL and CL. Unlike the penetrated helical structure of ClyA, the membrane inserted *β* barrel of AHL creates a more benign lipid-protein environment. The presence of the extracellular cap which interacts predominantly with the headgroups of the EL, does not induce significant leaflet heterogeneity despite an observed 9-fold increase in polar contacts which occur predominantly with the lipid headgroups, and unlike ClyA, to a much lesser extent with the alkyl chains of the lipid. As a consequence we did not observe significant differences in local order as revealed in the *g*(*r*), the lipid tilt angle and *S*_*C*−*C*_ across leaflets. Despite the large number of protein-lipid contacts with the EL, the extent of dynamic heterogeneity is reduced and the sub-diffusive regime is smaller when compared with ClyA.

### 4.2 Extent of Structural and Dynamic Perturbation

The quantification of the extent of structural and dynamic properties permits us to assess the relative changes between the two classes of proteins. A central finding in this manuscript, is that despite the large size, hydrophobic mismatch and uniquely different topologies and varying protein-lipid contacts (Fig. 6), perturbation to lipid order (in either leaflet) as reflected in the tilt angle and order parameter distributions (Figs. 3 and 4) as well as membrane thinning (Fig. 5), is short ranged, limited to ∼ 2.5 nm from the periphery of the either pore complex. Note that with periodic boundary conditions the average edge-to-edge distance from the pore complex for ClyA is about 7 nm and about 10 nm for AHL (ESI†). Substantially larger simulation box sizes would be required to assess the influence of periodic boundary conditions and study the region of influence around isolated pores. However, even in this relatively crowded regime of protein coverage, lipids with their inherent amphiphilic nature are able to restrict the protein influence to a small region around the pore complex, indicating the short ranged extent of the structural perturbation to lipids, commensurate with distances typically associated with van der Waals forces. This suggests that lipid-lipid interactions begin to dominate a few nm from the protein periphery to re-establish lipid packing and tail order thereby mitigating the influence of the inserted protein. This is also borne out by the relative invariance in the area-per-headgroup as a function of distance from the pore. On the other hand the spatial extent of perturbations to the lipid dynamics as revealed in the continuous survival probabilities (Fig. 10) and displacement maps (Fig. 11) extend outward to at least 4 nm in the case of ClyA and AHL.

Despite the differences in hydrophobic mismatch and structure of the toxins, lipid mobilities as revealed in the displacement maps lie in a similar range, further suggesting an inherent accommodation by the lipid bath. The large residence times of lipids (> 1 *µ*s) around both pore complexes CSP (Fig. 10) suggests a shell of lipids that is bound to the protein. This is similar to the notion of hydration shells associated with ions dissolved in water supporting the notion of an effective hydrodynamic radius which includes the shell of strongly bound lipids around the protein complex ^10^. Although the approach to the diffusive limit occurs at a shorter time window when compared with the bare membrane, the dynamic heterogeneity as revealed in the sub-diffusive exponents (Table 2) is similar to what is observed in a bare lipid membrane. Extended *µ*s long simulations would be required to further confirm these differences in the sub-diffusive and diffusive regions with the bare membrane.

### 4.3 Influence of Membrane Composition and Hydrophobic Mismatch

Since mammalian plasma membranes, which are biologically relevant in the context of bacterial toxins, are complex multicomponent systems consisting of hundreds of lipids, of which, phosphatidylcholine (PC) lipids comprise *>* 50% of the phospholipids, ^69^ PC lipids have been widely used to represent the plasma membrane in both simulations and experiments with PFTs. Although the membrane composition can influence the kinetics of pore formation, PFTs are notoriously membrane agnostic in their action and both ClyA and AHL have the ability to form pores in a wide range of membrane environments ranging from simple single component membranes, membranes with and without cholesterol and multicomponent membranes in erythrocytes which are commonly used to test lytic activity of proteins. Given that the pore formation for either ClyA or AHL is not impaired across a wide range of membrane compositions we restricted our study to DMPC an archetypical PC lipid. A molecular dynamics study on the hydrophobic mismatch ^70^ illustrated 17.6%, 13.9% and 15.1% mismatch for ClyA in 1,2-*dioleoyl-sn-glycero-3-phosphocholine* (DOPC):Cholesterol (70:30) DMPC and 1-*palmitoyl-2-oleoyl-sn-glycero-3-phosphocholine* (POPC):Cholesterol (70:30) membranes respectively. Thus across these different membrane compositions the changes were found to lie within a few percent. The choice of DMPC has interesting consequences in our study due to the differences in hydrophobic mismatch. The hydrophobic mismatch is greater in the case of ClyA when compared with AHL where the mismatch is minimal (Fig 5). In spite of these differences the spatial range of perturbations to both the lipid structure and residence times are similar for these disparate pore systems. While membrane compositions can influence the kinetics of pore formation, our main findings are expected to be qualitatively similar for simple membrane compositions typically used in experiments. ^19^ Specific differences in the local dynamics are likely to occur for example in the presence of cholesterol ^18,51^. STED-FCS experiments with ClyA in three component membranes show that the binding of ClyA to sphingomyelin and cholesterol rich domains reduces phase segregation in these raft forming membranes. ^21^ Molecular dynamics simulations of transmembrane proteins in multicomponent membranes reveal segregation of specific lipid types to the protein surface needed for specific functions of the transmembrane domains. ^25,71^ It would be interesting to study the local compositional changes in the case of assembled PFTs which do not have any specific function other than membrane perforation. Before concluding we point out that glass-like and heterogeneous dynamics which are observed in simple membranes with and without proteins are likely to be enhanced in the case of complex multicomponent membranes.

## 5 Summary and Conclusions

The study of the action of PFTs has largely focussed on mechanisms of pore formation, kinetic pathways and analysis of leakage and lysis kinetics combined with mutagenesis studies to unravel the influence of protein structure and role of specific residues on pore formation. The influence of the presence of toxin molecules and pores on the ensuing changes in lipid structure and dynamics has received less attention. Our MD simulations in a single component DMPC membrane reveals that relatively small sized PFT oligomers such as those formed by ClyA and AHL perturb the lipid structure and dynamics to within 4-5 nm from the pore complex, with short ranged structural perturbations restricted to within 2.5 nm commensurate with van der Waals type interactions. Thus the overall extent of perturbation is short ranged, typically involving several 100’s of lipids in the vicinity of the pore complex creating a shell of bound lipids that are likely to be intrinsically connected with protein mobility. ^10,11^ Thus lipids confer a conducive hydrophobic host membrane environment to assist spontaneous assembly and insertion of proteins, a process that is energetically favourable by design. The relatively local spatial extent of perturbation to membrane fluidity has implications on microvesicle shedding and budding events that are precursors to membrane repair processes designed to remove infected sites from the cell surface. ^6,7^

While MD simulations allow a detailed description of events at the molecular scale, several experimental methods can complement these findings. Recent super-resolution stimulated emission depletion coupled with fluorescence correlation spectroscopy (STED-FCS) spectroscopy on supported lipid bilayers have been used to study the inherent dynamic heterogeneity and non-Brownian dynamics associated with lipids in the vicinity of the PFTs. ^21,22^ Indeed, increased fluidity away from the pore complex has been observed due to cholesterol depletion with LLO (a cholesterol dependent cytolysin) binding, suggesting a more complex modulation mechanism in multicomponent membranes. ^18^ Advances in techniques such as neutron and X-ray scattering experiments can provide both lateral and in-plane variations in lipid order and dynamics at the atomic scale. ^72^ With improved instrumentation and single molecule methods probing dynamics at length scales currently studied in molecular dynamics studies is a distinct possibility. An understanding of the lytic action of PFTs and the concomitant lipid reorganization could potentially help in developing novel treatment protocols which seek to disrupt protein binding and pore formation ^73^, as well as understand neurodegenerative diseases such as Alzheimers which are caused by membrane-bound protein aggregates on the neuronal cell membranes, whose formation largely mimics self-assembly mechanisms followed by PFTs. ^74^

## Acknowledgements

RD and KGA thank Prof. Himanshu Khandelia for access to computational resources of the MEMPHYS group at the University of South Denmark. The authors thank the Department of Science and Technology (DST) for funding, the Supercomputer Education and Research Centre (SERC) for access to CrayXC40 and other high performance computing resources for post-processing simulation data and the Thematic Unit of Excellence in Computational Materials Science (TUE-CMS) as well for computational resources.

## Supplementary Information (ESI)

**Table S1:** System Details for ClyA and AHL Molecular Dynamics Simulations

|  | ClyA | AHL |
| --- | --- | --- |
| System size (number of atoms) | 526274 | 435739 |
| Simulation box size - L (nm) × W (nm) × H (nm) | $16.1 \times 16.1 \times 19.9$ | $17.3 \times 17.3 \times 14.4$ |
| Number of DMPC lipid molecules | 688 | 877 |
| Number of water molecules | 128832 | 99681 |
| Run-times of simulations (ns) | 500 | 500 |
| Salt concentration (mM) | 150 | 150 |
| Number of Na<sup>+</sup> ions | 623 | 449 |
| Number of Cl<sup>-</sup> ions | 539 | 456 |
| Temperature (K) | 310 | 310 |
| Pressure (bar, semi-isotropic) | 1 | 1 |

**Figure S1:**
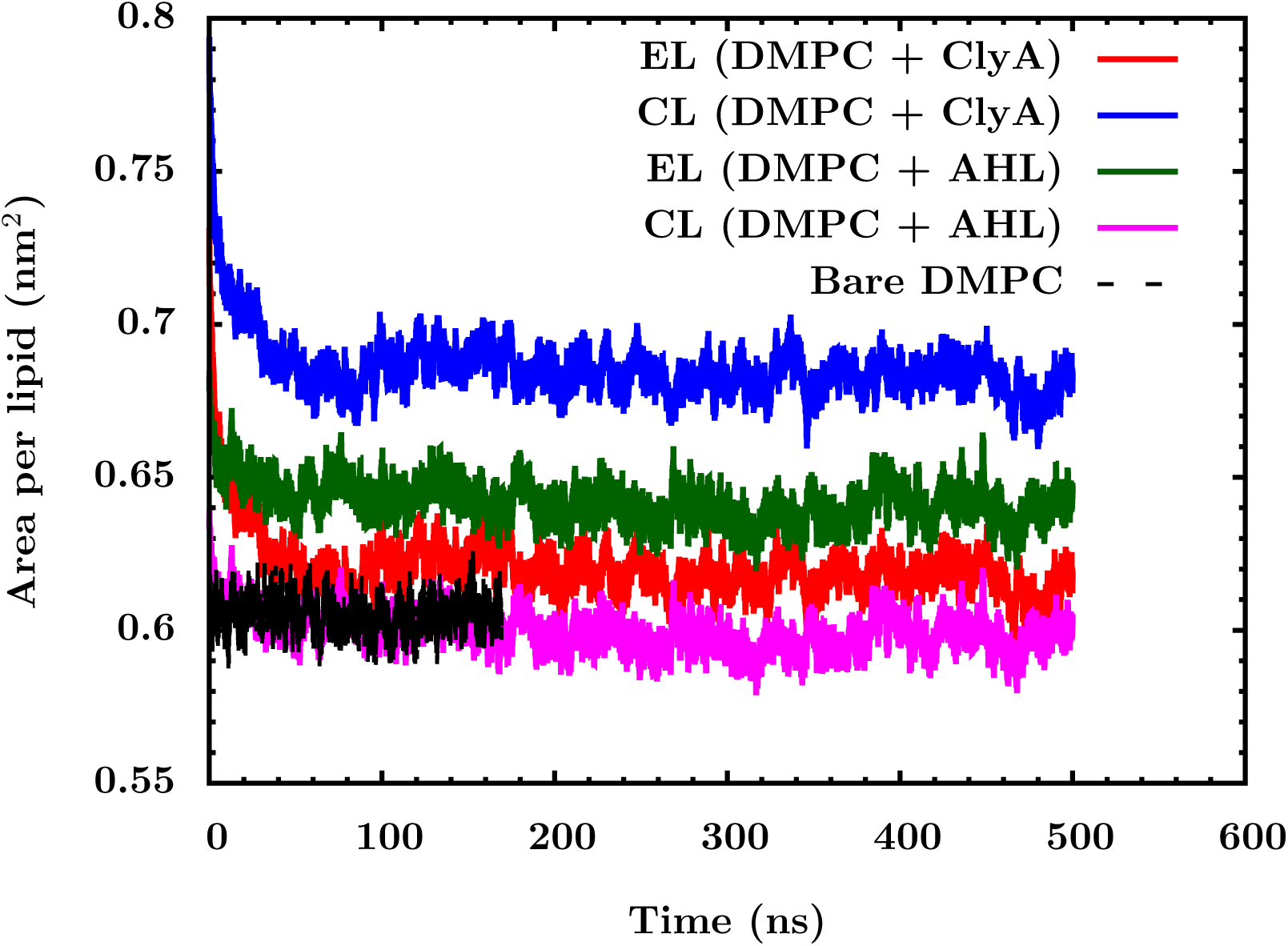
Area per lipid, *a*_*l*_, for the PFT bound DMPC bilayer. The values of *a*_*l*_ indicate that areal fluctuations stabilize within the first 100 ns.

**Figure S2:**
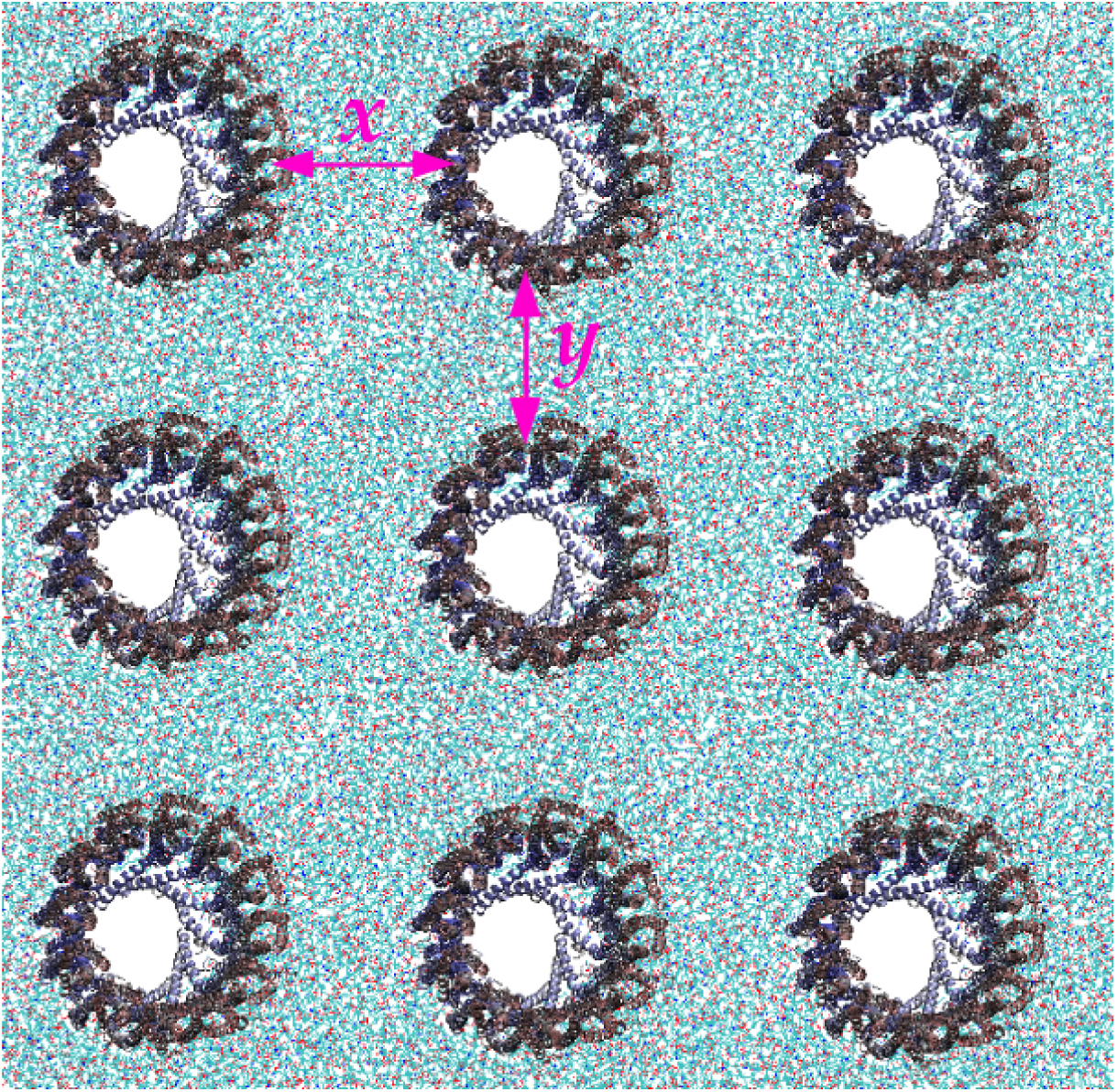
Periodic images of cytolysin A pores in the DMPC bilayer. The pores are separated at about 3.9 nm in the EL and 5.2 nm in the CL and are equidistant along the *x* and *y* directions.

**Figure S3:**
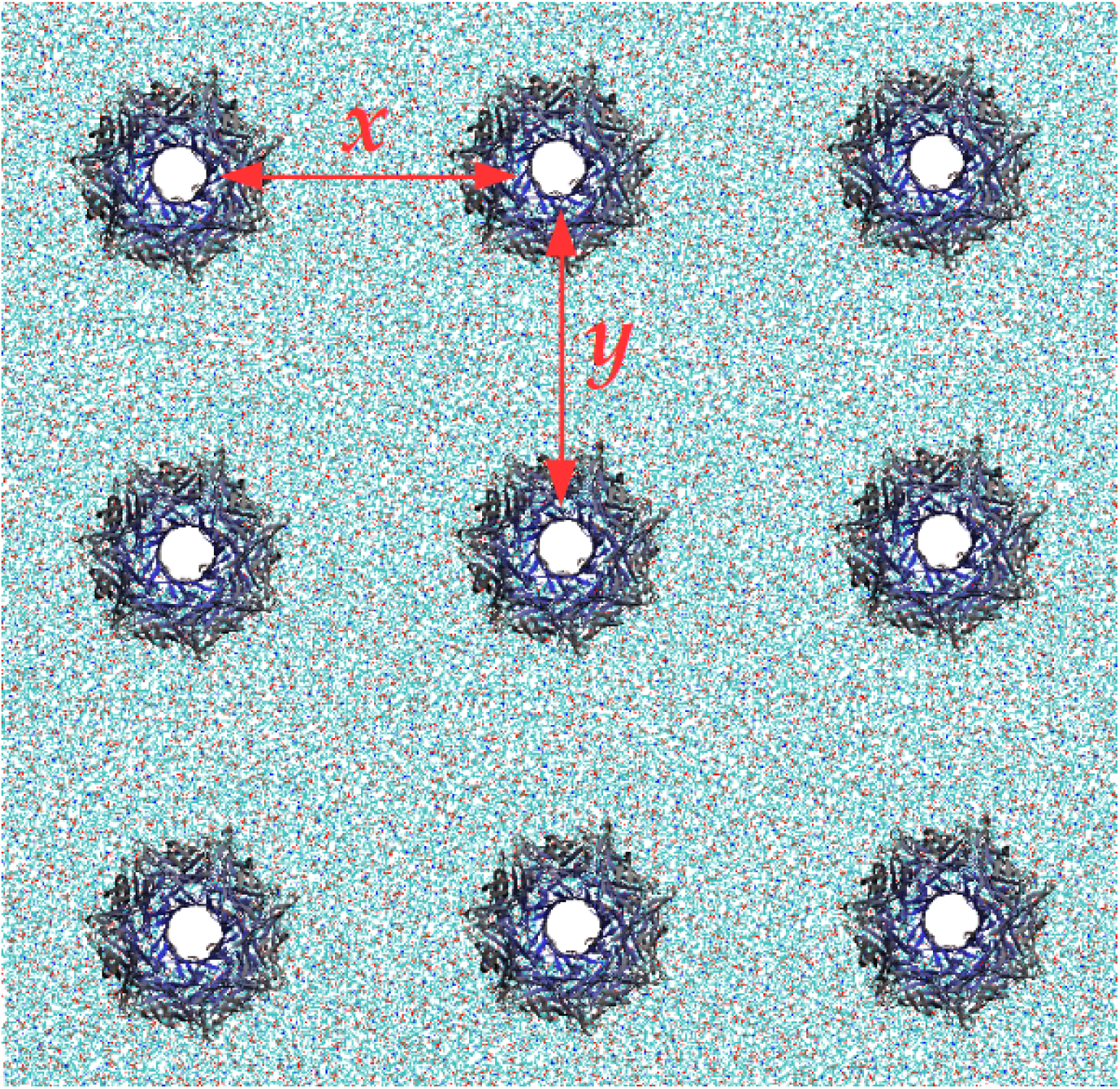
Periodic images of *α*-Hemolysin pores in the lipid bilayer. The pores are 6.4 nm away in the the EL and 6.8 nm away in the CL and are equidistant along the *x* and *y* directions.

**Figure S4:**
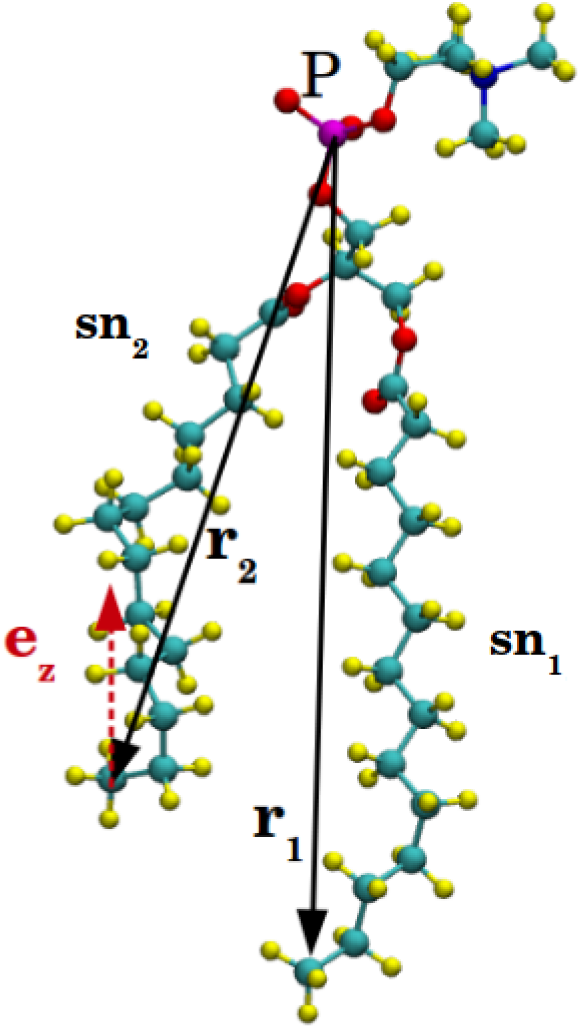
Illustration of the tilt angle subtended by the vector from the head group to the C-14 acyl carbon with the bilayer normal.

**Figure S5:**
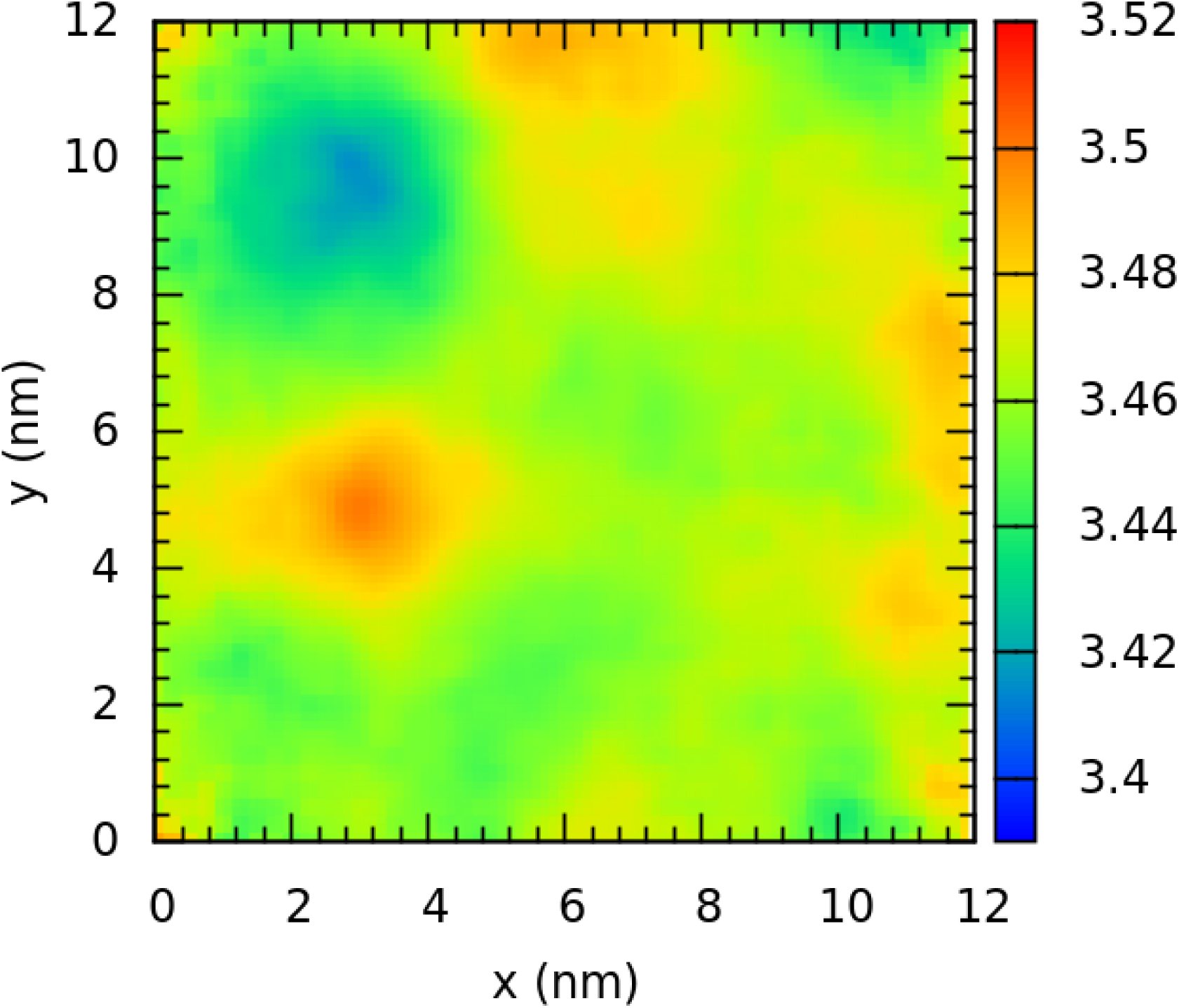
Thickness map of a bare DMPC bilayer at 310 K.

**Table S2:** Number of protein-membrane atomic contacts for the *α*-hemolsyin and cytolysin A pores. Values scaled to 100 with the maximum number of contacts are illustrated in Figures 11 & 13 of the main manuscript.

|  | AHL | ClyA |
| --- | --- | --- |
| Protein-CL | 25136.0 ± 999.4 | 85391.3 ± 2133.8 |
| Protein-CL (headgroup) | 6286.5 ± 480.1 | 15178.8 ± 965.6 |
| Protein-CL (sn1) | 11249.5 ± 689.9 | 36114.9 ± 1086.0 |
| Protein-CL (sn2) | 7600.0 ± 674.2 | 34097.6 ± 1520.1 |
| Protein-EL | 75643.0 ± 2055.8 | 75961.7 ± 1673.5 |
| Protein-EL (headgroup) | 44191.5 ± 1284.3 | 18828.7 ± 910.6 |
| Protein-EL (sn1) | 12667.0 ± 1047.5 | 25413.7 ± 1342.4 |
| Protein-EL (sn2) | 18784.4 ± 928.3 | 31719.3 ± 1443.6 |
| Polar AA-CL | 5170.8 ± 359.7 | 13310.0 ± 565.8 |
| Polar AA-CL (headgroup) | 2702.3 ± 283.1 | 2362.6 ± 401.8 |
| Polar AA-CL (sn1) | 1532.8 ± 228.8 | 6158.3 ± 350.4 |
| Polar AA-CL (sn2) | 935.7 ± 140.8 | 4789.1 ± 358.6 |
| Polar AA-EL | 37805.8 ± 1278.8 | 38686.3 ± 719.3 |
| Polar AA-EL (headgroup) | 25330.2 ± 885.2 | 13743.1 ± 612.8 |
| Polar AA-EL (sn1) | 4671.3 ± 523.8 | 10471.6 ± 612.4 |
| Polar AA-EL (sn2) | 7804.3 ± 441.3 | 14471.7 ± 655.9 |
| Non-polar AA-CL | 19965.2 ± 783.3 | 72081.3 ± 1773.6 |
| Non-polar AA-CL (headgroup) | 3584.2 ± 338.8 | 12816.1 ± 761.1 |
| Non-polar AA-CL (sn1) | 9716.6 ± 552.4 | 29956.6 ± 1024.0 |
| Non-polar AA-CL (sn2) | 6664.3 ± 574.7 | 29308.5 ± 1315.1 |
| Non-polar AA-EL | 37837.2 ± 976.6 | 37275.4 ± 1326.4 |
| Non-polar AA-EL (headgroup) | 18861.3 ± 521.3 | 5085.6 ± 399.8 |
| Non-polar AA-EL (sn1) | 7995.7 ± 694.7 | 14942.1 ± 967.9 |
| Non-polar AA-EL (sn2) | 10980.1 ± 663.1 | 17247.6 ± 943.0 |
| RKWY-CL (headgroup) | 828.1 ± 180.1 | 626.5 ± 129.9 |
| RKWY-CL (sn1 & sn2) | 76.0 ± 32.3 | 8258.6 ± 365.6 |
| RKWY-EL (headgroup) | 17048.5 ± 534.0 | 9261.4 ± 405.6 |
| RKWY-EL (sn1 & sn2) | 4697.8 ± 216.1 | 19612.9 ± 607.9 |

